# Fetal cerebellar development: 3D morphometric analysis of fetal postmortem MRI among Chinese fetuses

**DOI:** 10.1101/2021.10.10.462694

**Authors:** Emmanuel Suluba, Liu Shuwei

## Abstract

The development of the cerebellum starts from early gestational period and extends postnatal. Because of its protracted development, the cerebellum is susceptible to developmental anomalies such as Dandy-Walker malformations, Blake’s pouch cysts and vermin hypoplasia. Measurements of fetal cerebellar parameters of a normal growing fetus in each week of gestation is important for setting up morphometric standards and hence used as clinical reference data. Any deviation from the normal cerebellar parameters alerts the clinicians for the possibility of presence of cerebellar malformations.

**Study objective:** The objective of this study was to assess the fetal cerebellar growth by quantifying the following parameters; fetal cerebellar volume, anterior-posterior diameter and superior-inferior diameter.

**Methods:** We used 3T and 7T MRI to scan the postmortem fetal brains at different stages of development and subsequently analyze the images using ITK-SNAP software.

**Results:** The mean superior-inferior cerebellar diameter was found to be 19.12±2. 70mm.The linear(y=b_o_+b_1_t) model was the best fit (r^2^=0.996, F=32022.961) to describe the relationship between the gestational age and the superior-inferior diameter(y=5.89+0.49t). There was significant correlation between the superior-inferior cerebellar diameter and the gestation age, Pearson correlation coefficient of 0.999, r=0.001. The median cerebellar volume was 8607.7mm, the mean rank high among males(78.12) as compared to female(68.25).There was no statistically significant difference of the cerebellar volume between males and females (u=2193.5,p=0.16).The quadratic(y=b_o_+b_1_t+b_2_t^2^) model was the best fit regression equation (r^2^=0.994,F=10791.157) describing the relationship between the cerebellar volume and the gestational age. The median anterior- posterior diameter was 12.45 mm. There was significant correlation between anterior-posterior diameter and the gestational age with Spearman’s rho of (0.997, p=0.01). The linear model was the best fit the best fit model (y=b_o_+b_1_t) describing the relationship between anterior-posterior diameter and the gestational age(y=3.31+0.5t) r^2^=0.998, F=70646.838

**Conclusion:** Significant correlation between the superior-inferior cerebellar diameters, the anterior-posterior cerebellar diameter and the gestational age was found. These two linear parameters follow the first-degree polynomial in relation to the gestational age. The cerebellar volume follows the second-degree polynomial as it increases with the gestational age and correlate significantly with the gestational age. This study has provided new insight to the development of the cerebellum, and setup a benchmark data of which the deviation from it will alert the clinicians for the possibility of presence of cerebellar malformations.

## Introduction

The development of human cerebellum starts from the early embryonic period and continue one year after birth. This long period of development(protracted development) makes the cerebellum vulnerable to common developmental disorders such as Dandy-Walker malformations and pontocerebellar hypoplasia(Ten Donkelaar and Lammens 2009).There are four important phases of cerebellar development namely, cerebellar territory characterization; the formation of ventricular zone and rhombic lip forming Purkinje cells, deep cerebellar nucleus and granule cells; the migration of granule layer from external to internal and lastly the cerebellar circuitry formation (ten Donkelaar et al. 2003).Apart from the formation of cerebellar hemispheres, the cerebellar vermis, a small, hindbrain structure develop from the rostral segments of the rhombencephalon in early gestation(Martinez et al. 2013, Millen and Gleeson 2008).The cerebellum contribute to both motor and non-motor functions such as movement, proprioception, and cognition(Basson and Wingate 2013, Koziol et al. 2014, Ben-Yehudah, Guediche, and Fiez 2007). Cerebellar malformations are common (Santoro et al. 2019, Howley et al. 2018) and have long term consequences in children(Pinchefsky et al. 2019, Abel and Tahir 2019),therefore frontiers in obstetric and radiological societies emphasize the important for routine screening of the cerebellum and vermis in utero(Haratz et al. 2019, Quarello et al. 2014). Early detection (prenatal diagnosis of the cerebellar abnormality is essential and have some prognostic impacts (Patek et al. 2012, Gandolfi Colleoni et al. 2012, Wuest et al. 2017).Obstetric ultrasound has been a routine diagnostic tools in evaluation of fetal well-being in utero(Whitworth, Bricker, and Mullan 2015, Henrichs et al. 2019), however, the fetal cerebellar structures such as the vermis, being small in size and deep in skull base, the routine axial sonography may not suffice to depict the subtle changes during cerebellar development(Yang et al. 2012, Pugash et al. 2008). Recently the improvement of fetal ultrasound techniques have increased the diagnostic accuracy(Tonni, Grisolia, and Sepulveda 2014, Bertucci et al. 2011, Zalel et al. 2009).However, the use of fetal MRI provides additional information that can eventually influence the diagnosis the line of clinical management(Imamoglu et al. 2013, Porat et al. 2002). Recently the development of neonatal imaging coil has improved the signal-noise ratio and therefore provide the good quality images(Barkovich 2006).MRI can also predict the fetal later neurodevelopment (Anderson, Cheong, and Thompson 2015).In this study, we are compelled to investigate the changes in cerebellar volume, anterior-posterior and superior-inferior diameters of the normal developing fetus in order to setup normal morphometric standards and subsequently be able to detect the deviation from the normal growing cerebellum.

## Material and Methods

### Fetal specimens and Image Acquisition

The consent for use of fetal specimens was obtained from both parents and the approval to conduct the study was provided by ethical committee of Shandong University School of Medicine. We acquire the specimens after medically indicated abortions. Medical conditions such as severe intrauterine growth restriction secondary to pregnancy induced hypertension, congenital fetal malformations unrelated to the brain, severe maternal infections and intrauterine fetal death due to traumatic conditions were the reasons for either spontaneous or medically induced abortions.

We performed 3T and 7T MR imaging for clear images and accurate 3D reconstruction. We used both 3T MR and 7T MR imaging to scan the fetuses at 18-37weeks. We obtained clear images at every stage of development (Images 1a, 1b, and 2,3,4,6 and7). Postmortem MRI imaging ensures that the measurement we obtained are accurate, it mitigates the motion artifacts and there are literally no field strength limitations. We used the following scan protocols; 7T micro-MR imaging (70/16 Pharma Scan; Bruker Bio spin, Ettlingen, Germany). We scanned fetuses aged 18-22 weeks by using the rat body coil inner diameter 60mm. The acquisition parameters of T1-weighted images were the following: TR/TE, 384.4/15.8 MS; matrix size, 512 _ 512; slice thickness, 0.8 mm; number of excitations, 1; FOV, 6_6 cm; voxel size, 0.8 _ 0.12 _ 0.12 mm3. The acquisition parameters of T2-weighted slice images were the following: TR/TE, 17,000/50 MS; matrix size, 256 _ 256; slice thickness, 0.5 mm; number of excitations,4; FOV, 6 _ 6 cm; voxel size, 0.5 _ 0.23 _ 0.23 mm3.Due to the coil limitations, the specimen aged specimens at 23–37 weeks of gestation were scanned with a Signa 3T MR imaging scanner (Healthcare, Milwaukee, Wisconsin). The acquisition parameters of T1-weighted slice images were the following: TR/TE, 2580.0/23.4 MS; matrix size, 512_512; slice thickness, 2 mm; number of excitations, 1; voxel size, 2 _ 1.9 _ 1.9 mm3. The acquisition parameters of T2-weighted slice images were the following: TR/TE, 4600.0/111.6 ms; matrix size, 512 _ 512; slice thickness, 2mm; number of excitations, 1; voxel size, 2 _ 1.9 _ 1.9 mm3.

### Image Processing and data Measurement

A cross-sectional evaluation of fetal brains images to obtain the cerebellar measurements namely, the cerebellar volume, anterior- posterior diameters (APD) and the inferior-superior diameters (SID) was done. Images were uploaded and all three image volumes were investigated by two experienced radiologists using ITK-SNAP software (Yushkevich, Gao, and Gerig 2016, Yushkevich et al. 2019) All the measurements were done by two individuals for quality assurance and Kappa values were calculated for inter-observer variability. The volume of the cerebellum was automatically calculated after manual segmentation followed by 3D reconstruction (Image 5). The linear measurements (the anterior posterior diameters and the inferior-superior diameters were measured). The cerebellum was studied in all three planes The SID was measured by referring on the median sagittal plane of the fetal brain and was taken as the distance between the upper and lower margins of the cerebellum. The APD of cerebellar vermis was measured by referring to the median sagittal plane of the fetal brain and was taken as the distance between from the posterior wall of the fourth ventricle to the most dorsal margin of the cerebellum.

### Statistical analysis

We used SPSS software version 23 and GraphPad Prism 9.2 for data entry, cleaning and analysis. We tested the data for normality using Shapiro-Wilk, Kolmogorov – Smirnov test and Quantile-Quantile plots. Non-parametric test, Mann-Whitney U test and Spearman-ranking correlation coefficient were used for data which were not normally distributed in order to establish the differences between sexes and the correlation respectively. The data were expressed as median as the measure of central tendency. Parametric test, the independent T-test and Pearson correlation coefficient were used for normal distributed data. The data were expressed as the mean and standard deviation. We compared the cerebellar volume between sexes by Mann-Whitney test. The correlation between the cerebellar volume and the gestation age was established and Spearman-ranking correlation coefficient was reported with is its level of significance. We established the best fit regression equations and several mathematical models were tested to establish the best fit model. The P-value of < 0.005 indicated statistically significant difference. The inter-observer variability was established; Cohen’s Kappa value was calculated and reported with its level of significance. We also calculated the 95% confidence interval for Kappa and the obtained value was interpreted (Warrens 2014, Hsu and Field 2003). In addition, we plotted the Bland-Altman plots for inter-observer differences. Several mathematical models were tested to find out the best fit model.

## Results

A total of 144 fetuses were included in this study. The gestational age was between 18 to 37 weeks, with a mean age 27.5±5.8 weeks. Gestational age distribution is shown in Figure 1. The females were 86 while males were 62.The median cerebellar volume was at 8607.7mm.The mean rank was high among males(78.12) as compared to female(68.25).There was no statistically significant difference of the cerebellar volume between males and females (u=2193.5,p=0.16).The quadratic(y=b_o_+b_1_t+b_2_t^2^) model (Fig.3 and Fig.5)was the best fit regression equation with the largest r^2^=0.994,F=10791.157.The was significant correlation between the cerebellar volume and the gestational age with Spearman’s rho of (0.997,p=0.001).To establish the inter-observer variability Cohen’s Kappa was calculated; Kappa=0.007(p<0.001),95%CI(−0.005,0.018),Therefore there was slightly agreement between the two observers in the measurements of cerebellar volume. We plotted the Bland Altman plot using GraphPad prism (9.2.0) (Fig.17) for inter-observer variability. The 25, 75 and 95 cerebellar volume percentiles were calculated as in shown in (Fig.6) and (Table 1). Also, the comparison between male and females are shown in box plots (Fig.2). The cerebellar volume was increasing with the gestation age as shown in the scatter diagrams (Fig.4)

**Figure1:**
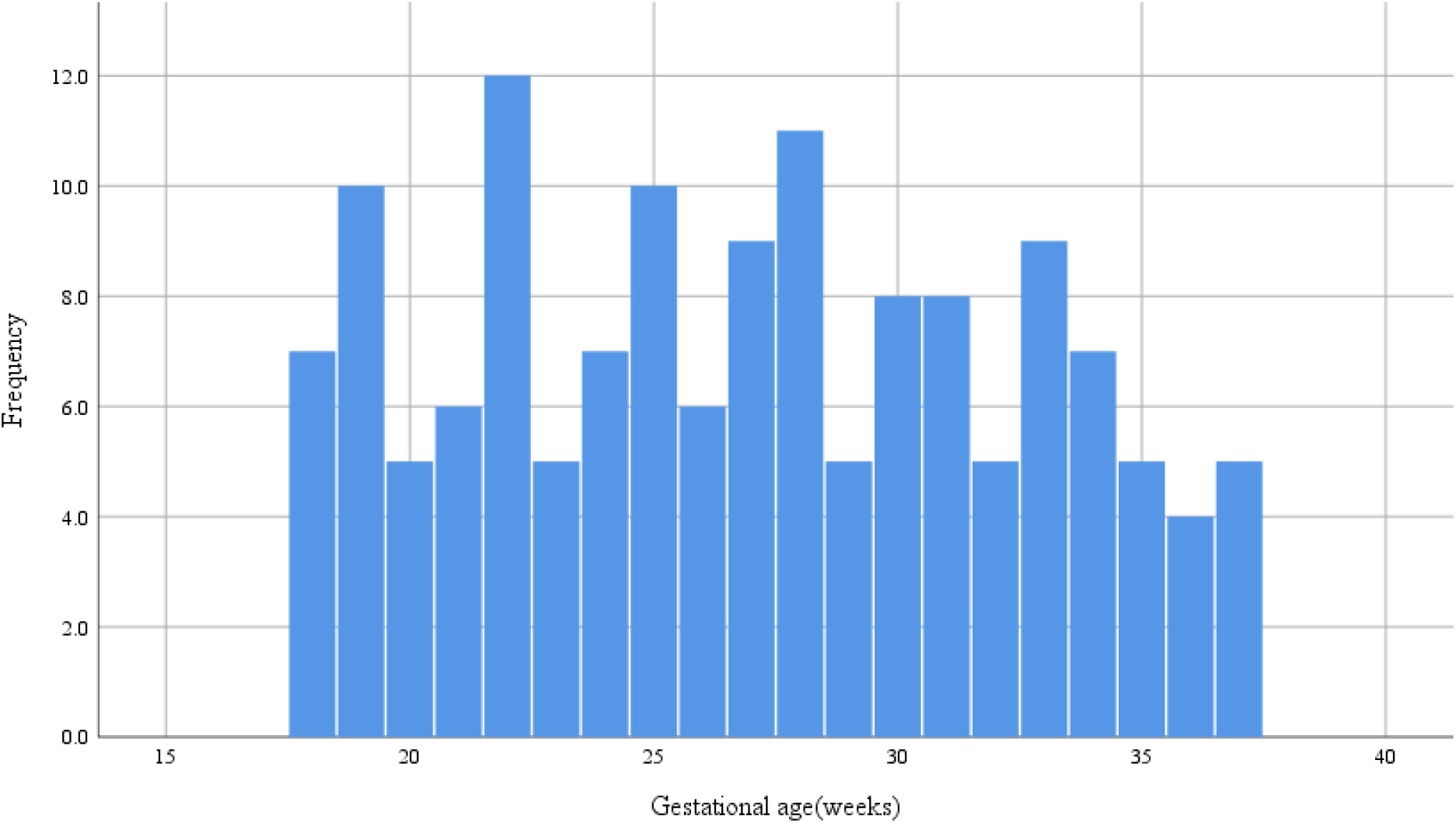
The Histogram shows the number of fetuses and gestational age distribution

**Figure 2.**
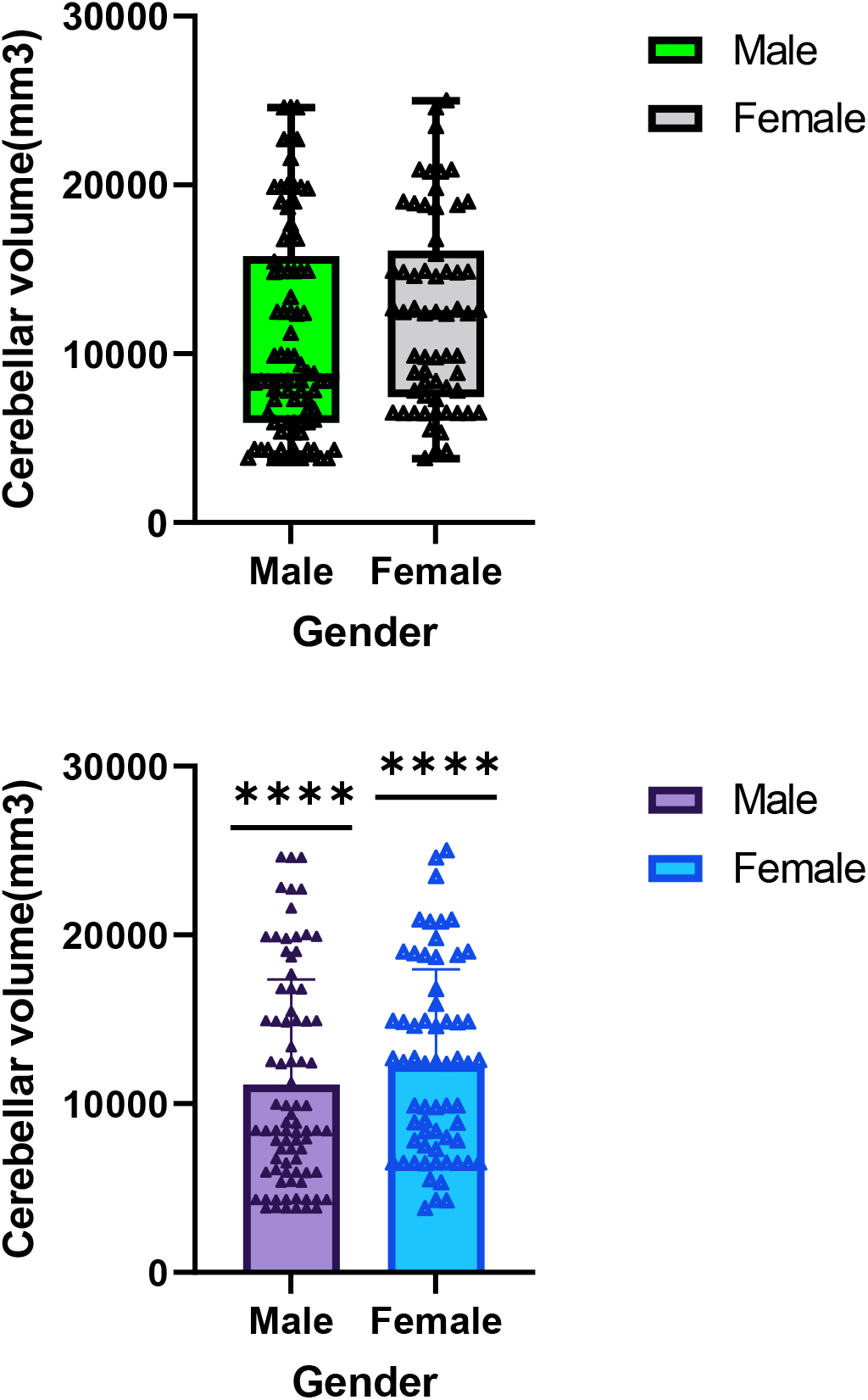
Box plot and median error pairwise comparison shows the difference in cerebellar volume between males and females

**Figure 3:**
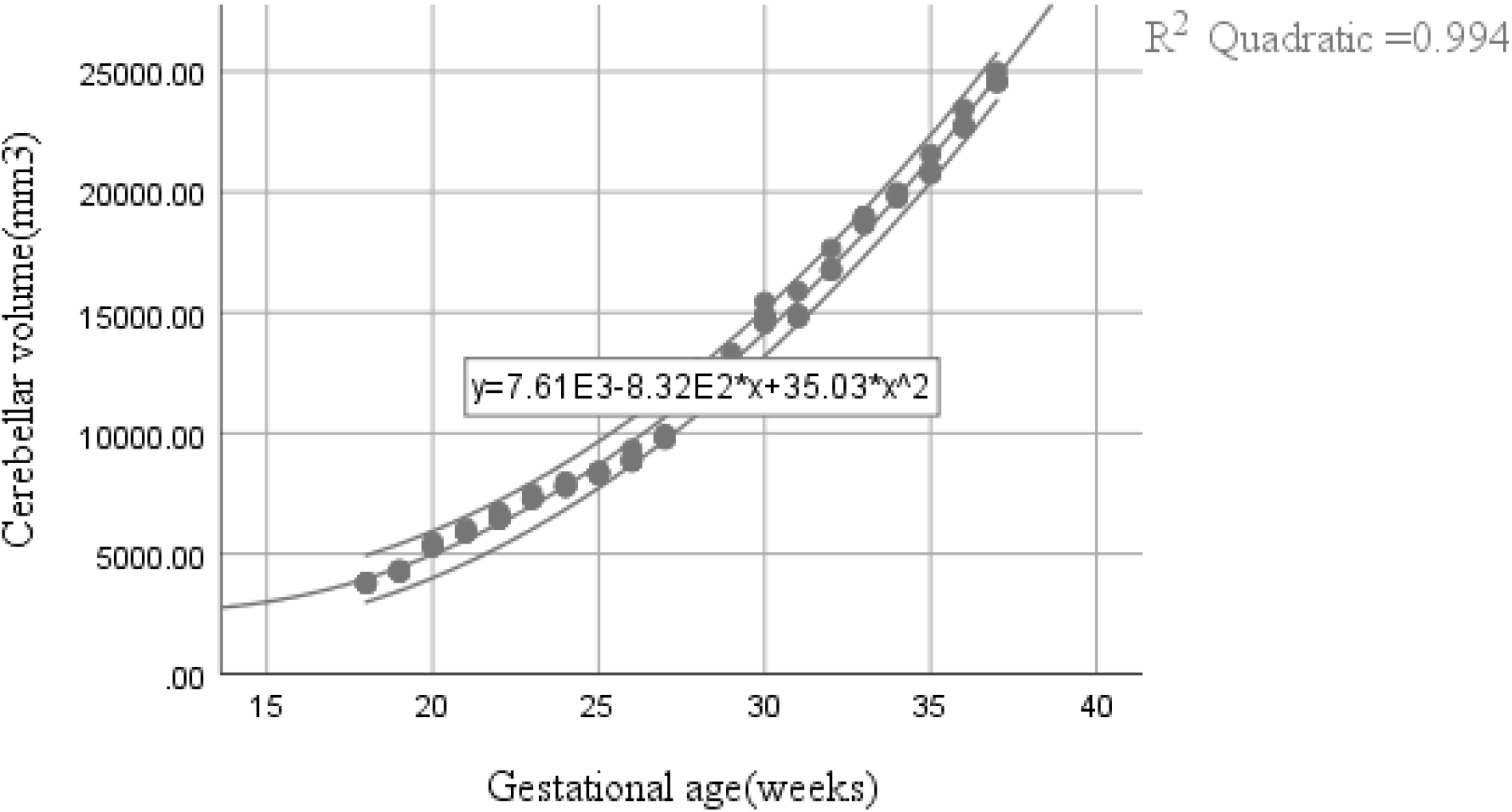
The cerebellar volume and the gestational age, the second degree-polynomial with 95%CI

**Figure 4:**
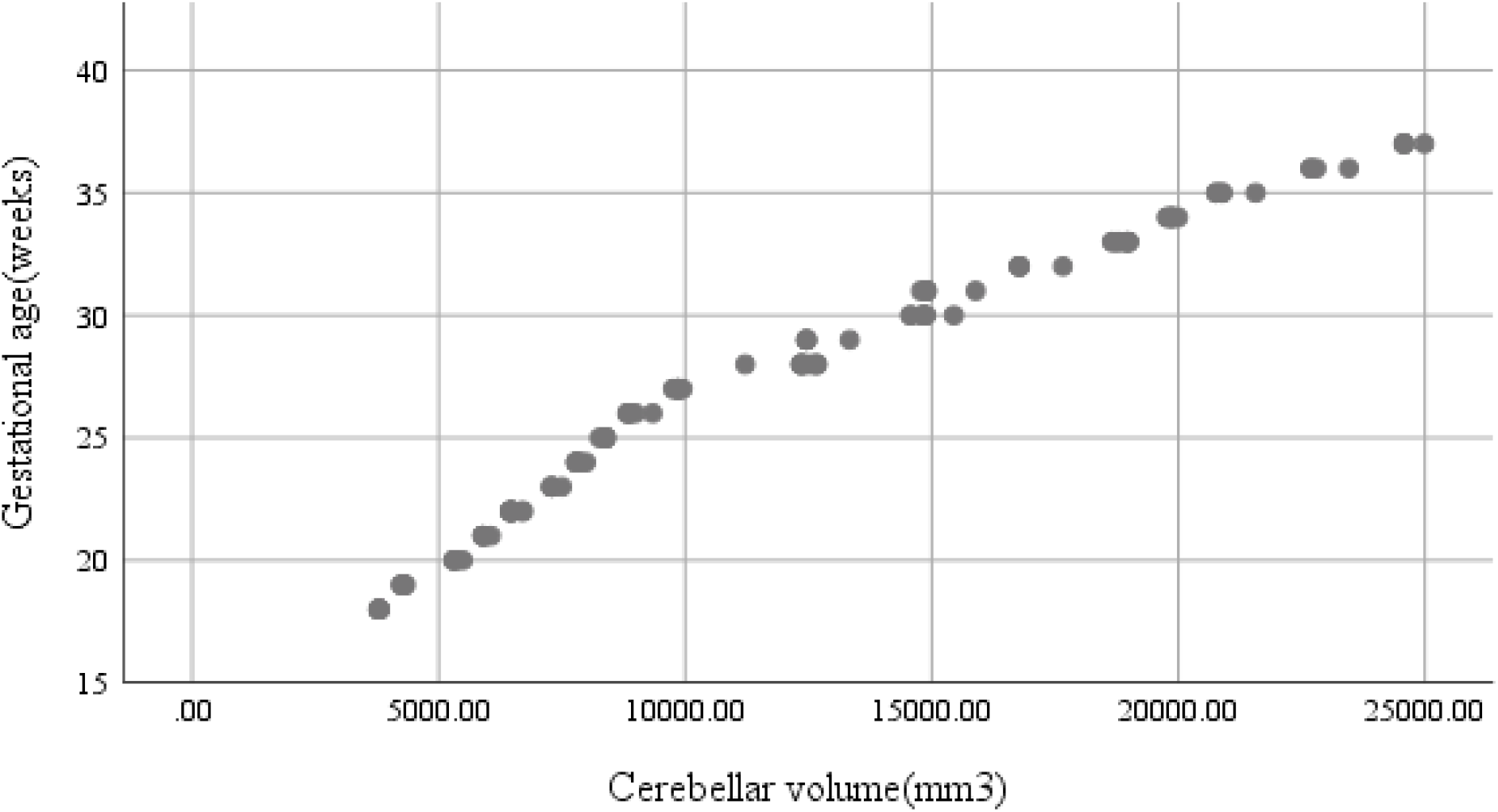
Scatter plot, cerebellar volume and the gestational age

**Figure 5:**
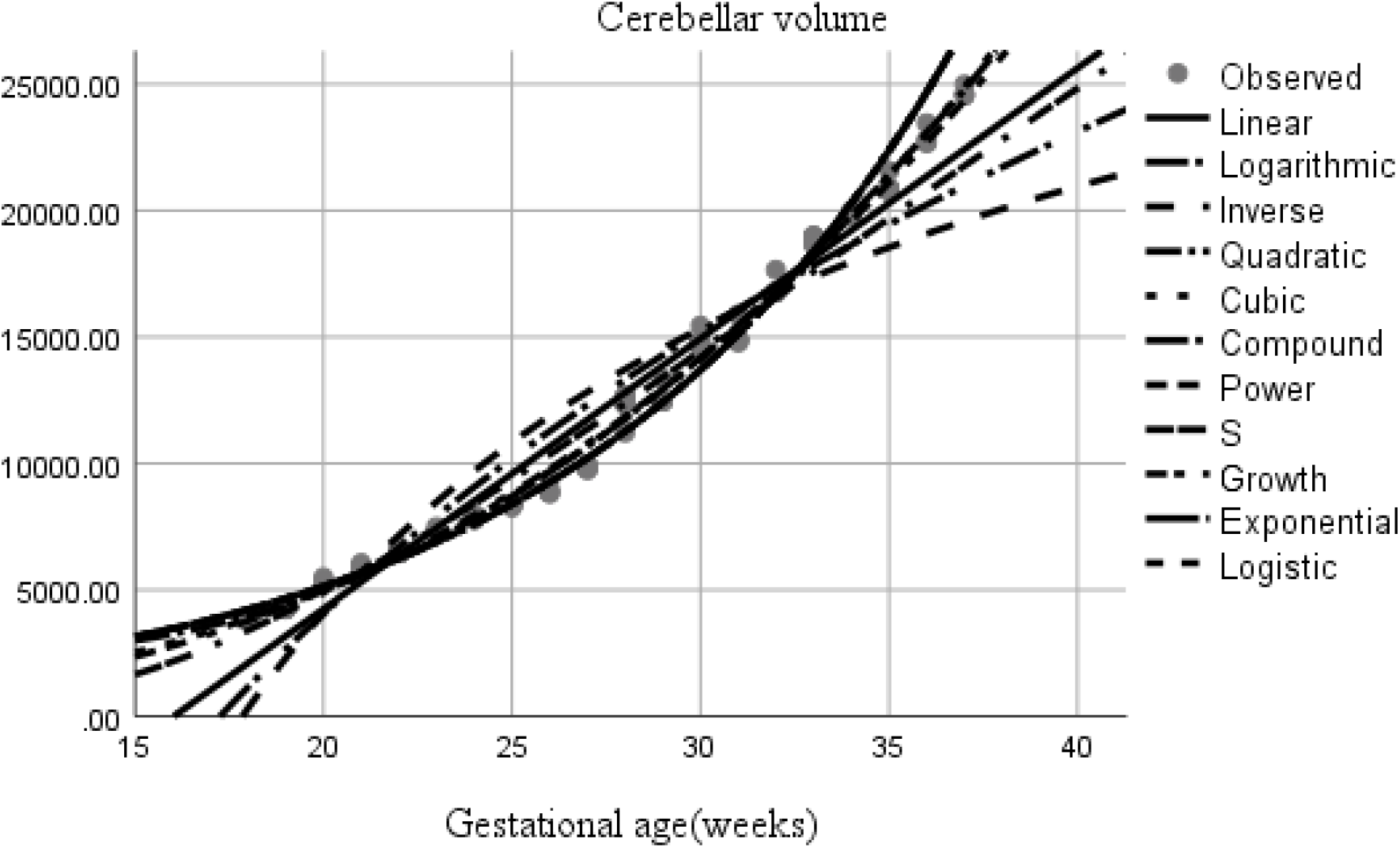
Shows several mathematical models tested, quadratic being the best fit regression model.

**Figure 6:**
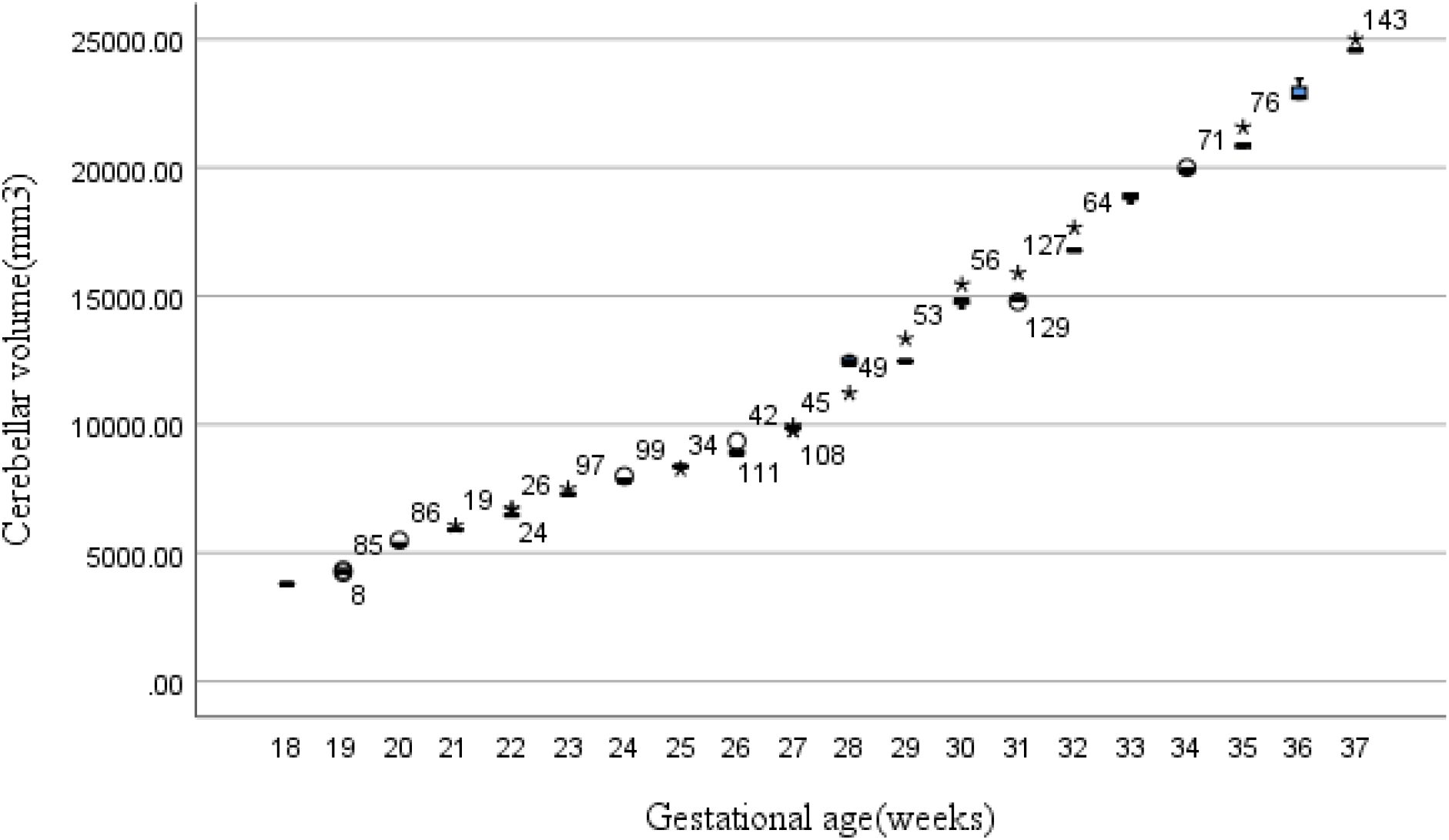
Trends of cerebellar volume and gestational age

**Table 1:** Cerebellar volume percentiles

| Gestational age(weeks) | 25 | 50 | 75 |
| --- | --- | --- | --- |
| 18 | 3780.7000 | 3792.6000 | 3795.2000 |
| 19 | 4265.7000 | 4280.7000 | 4291.9500 |
| 20 | 5300.3500 | 5320.4000 | 5422.3500 |
| 21 | 5897.0000 | 5900.4000 | 5954.1250 |
| 22 | 6456.0750 | 6471.1500 | 6487.8000 |
| 23 | 7286.5500 | 7288.5000 | 7393.8500 |
| 24 | 7787.4000 | 7799.2000 | 7887.3000 |
| 25 | 8353.7500 | 8366.6000 | 8379.8500 |
| 26 | 8832.0000 | 8860.3500 | 9076.3250 |
| 27 | 9815.4500 | 9860.6000 | 9870.4000 |
| 28 | 12346.4000 | 12376.6000 | 12646.7000 |
| 29 | 12455.8500 | 12456.9000 | 12901.5850 |
| 30 | 14639.6750 | 14828.6000 | 14874.4250 |
| 31 | 14859.6000 | 14887.8000 | 14897.5000 |
| 32 | 16767.9000 | 16776.9000 | 17222.5000 |
| 33 | 18724.9000 | 18897.4000 | 18980.6500 |
| 34 | 19779.4000 | 19867.5000 | 19877.8000 |
| 35 | 20767.9000 | 20887.4000 | 21227.9000 |
| 36 | 22680.7000 | 22743.2000 | 23292.1500 |
| 37 | 24568.3000 | 24568.6000 | 24787.8500 |

The mean superior-inferior cerebellar diameter was 19.12±2.70mm.The linear(y=b_o_+b_1_t) model was the best fit (Fig 8 and Fig.11) with the largest r^2^=0.996,F=32022.961) presenting the relationship between the gestational age and the superior-inferior diameter(y=5.89+0.49t).There was significant correlation between the superior-inferior cerebellar diameter and the gestation age, the Pearson correlation coefficient p= 0.999,r=0.001. There was no significant difference between males and females for the superior-inferior cerebellar diameter as Levene’s test for equality of means yielded t (142) = −1.399, p=0.048, equal variances assumed. The inter-observer variability Kappa=0.07(p<0.001),95% CI (0.028,0.11), there was slightly agreement between the two observers in the measurements of inferior-superior diameters as per Bland-Altman plots(Figure 18) The 25, 75 and 95 superior-inferior cerebellar diameter percentiles were calculated (Fig.10) and (Table 2). The superior-inferior cerebellar diameter was increasing with the gestation age as shown in the scatter diagrams (Fig.7). The comparison between male and females are shown in box plots (Fig.9)

**Figure7.**
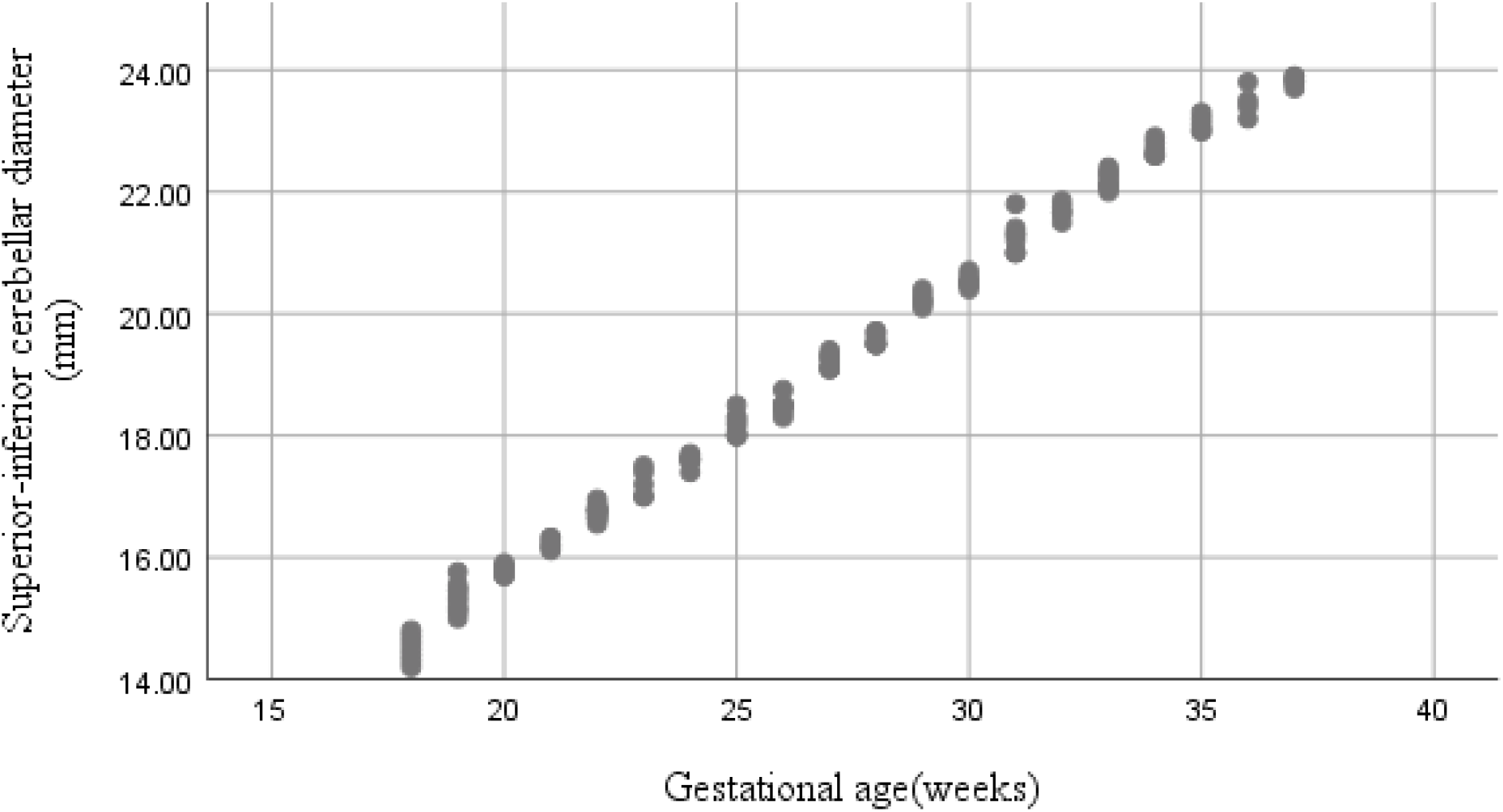
Scatter plot, superior-inferior and the gestational age

**Figure 8:**
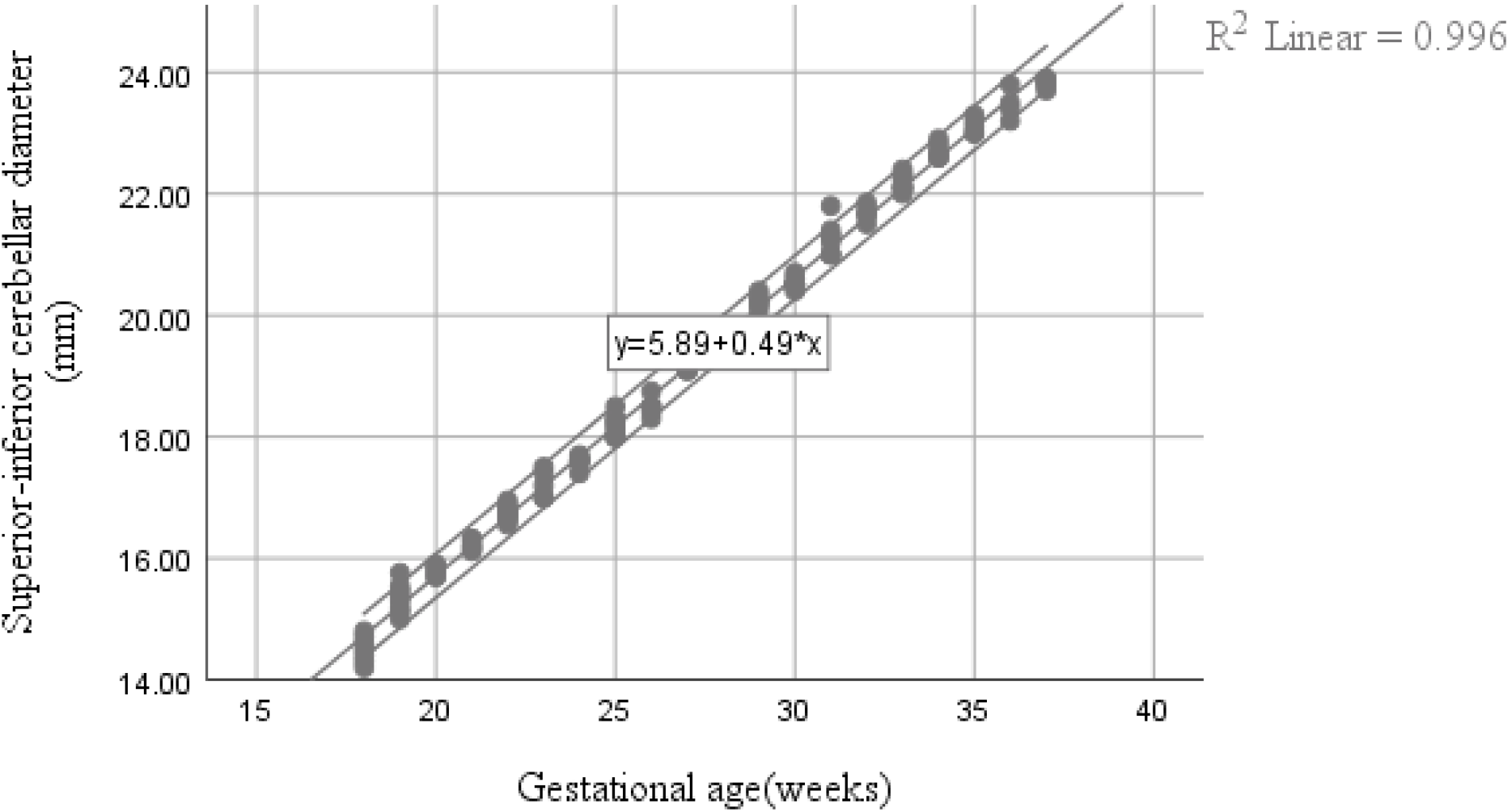
The superior-inferior diameter and the gestational age, linear function with 95%CI

**Figure 9.**
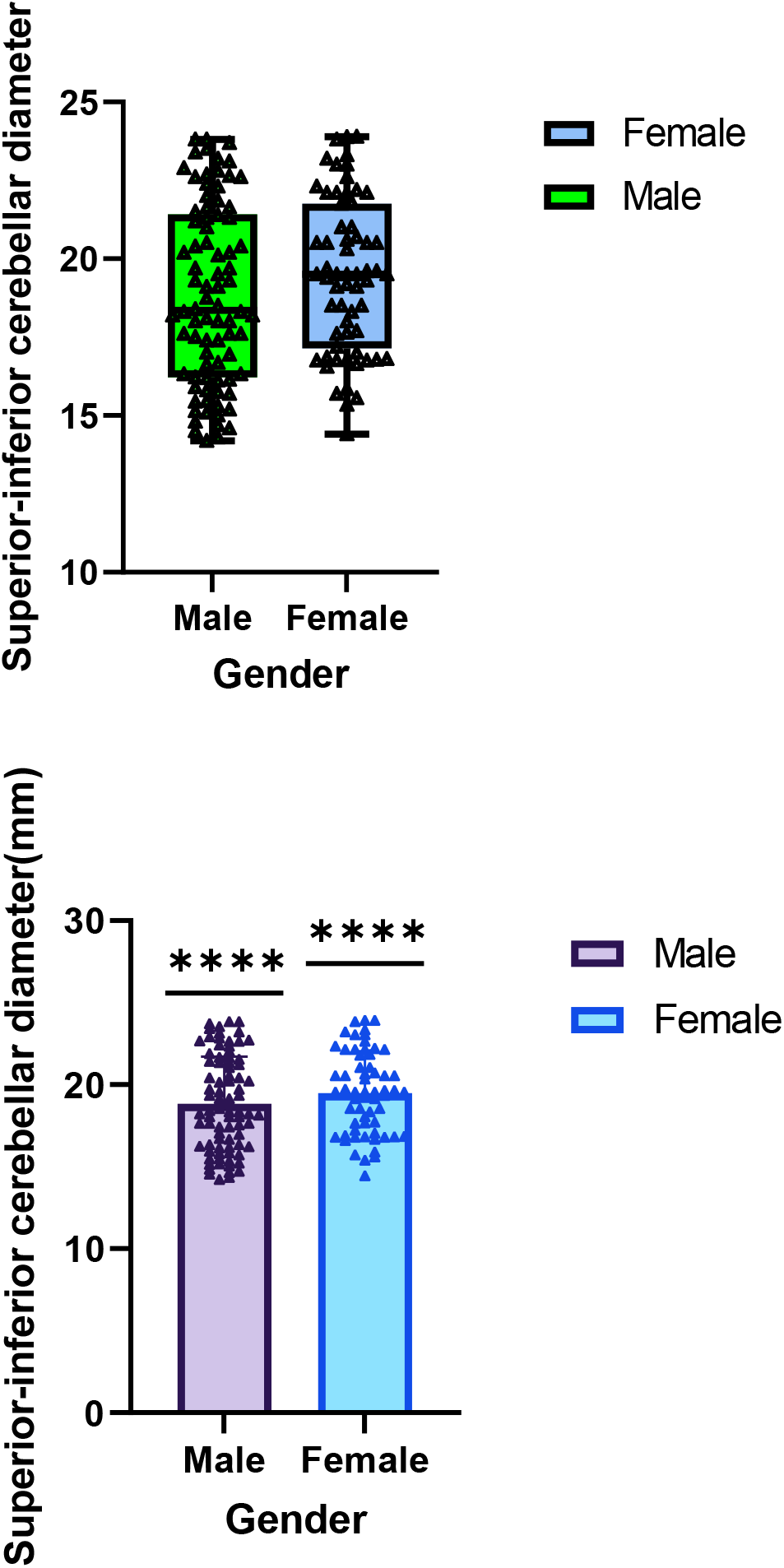
Box plot and median error pairwise comparison shows the difference in superior-inferior diameter between males and females

**Figure 10.**
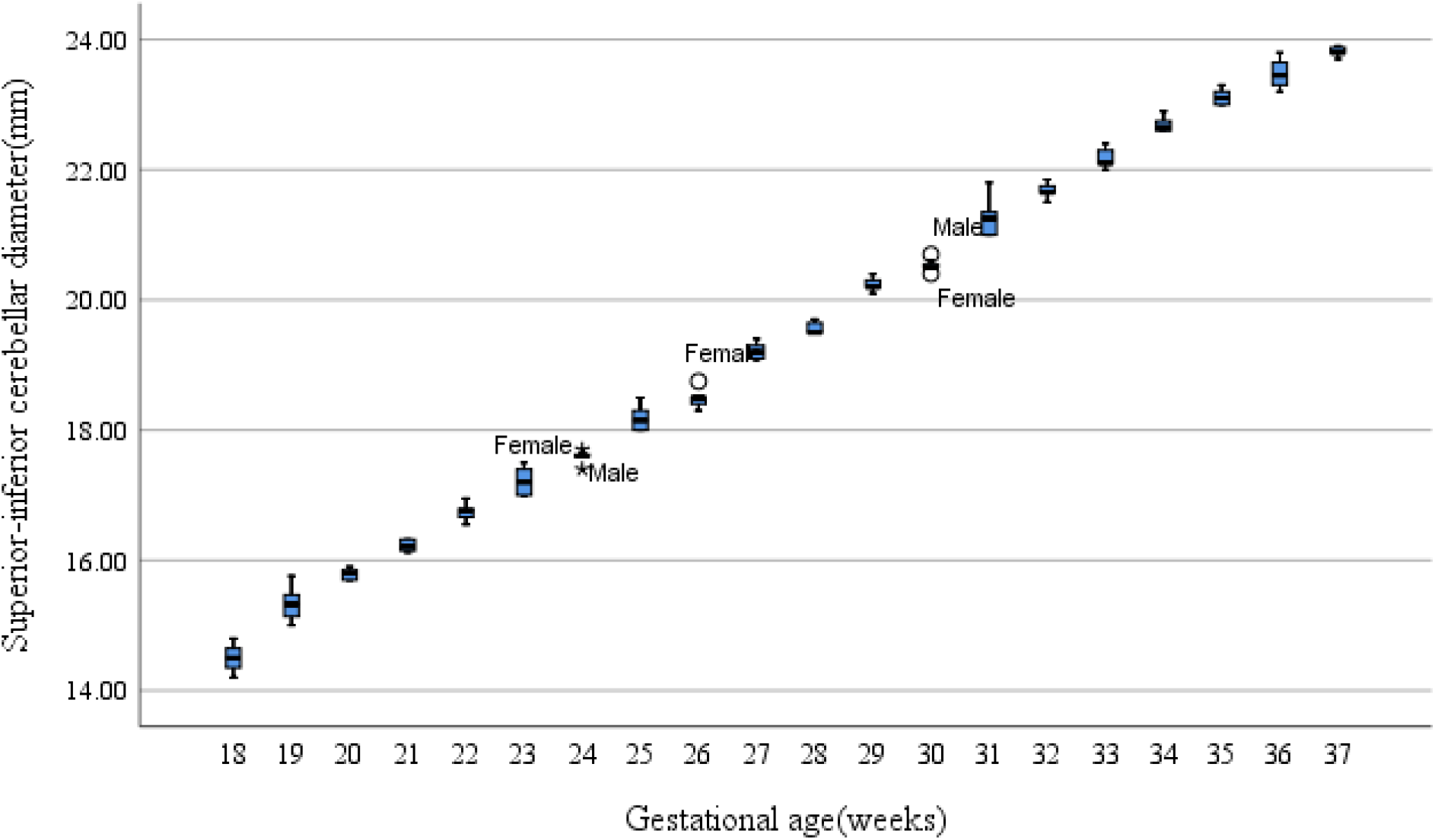
Trends of superior-inferior cerebellar diameters and gestational age

**Figure 11.**
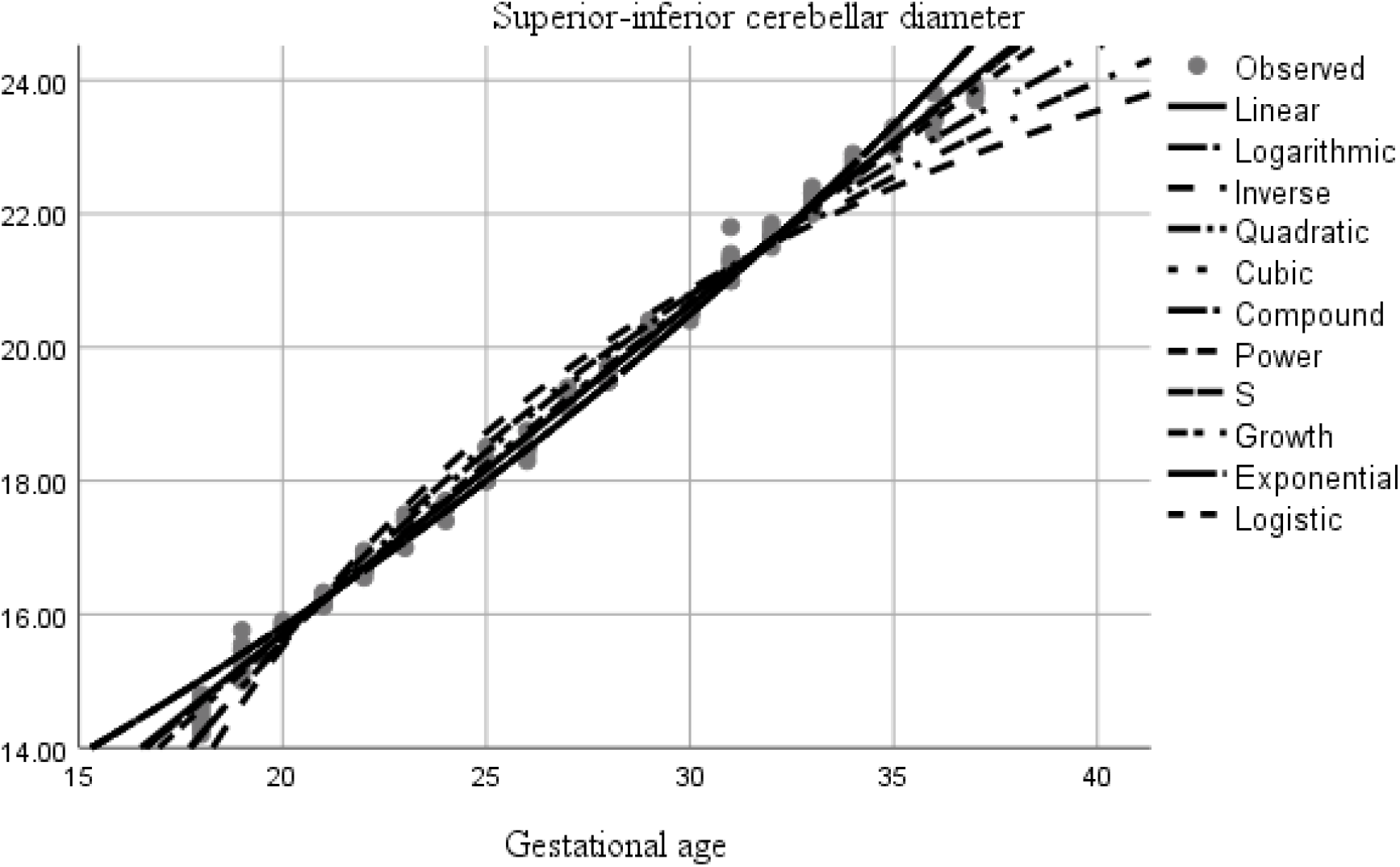
Shows several mathematical models tested, the linear model being the best fit regression model

**Table 2.** Superior-inferior cerebellar diameter percentiles

| Gestational<br>age(weeks) | 25 | 50 | 75 |
| --- | --- | --- | --- |
| 18 | 14.3000 | 14.5000 | 14.7000 |
| 19 | 15.1300 | 15.3250 | 15.4900 |
| 20 | 15.7000 | 15.8000 | 15.8750 |
| 21 | 16.1425 | 16.2100 | 16.3200 |
| 22 | 16.6575 | 16.7600 | 16.7950 |
| 23 | 17.0000 | 17.2000 | 17.4500 |
| 24 | 17.6000 | 17.6000 | 17.6500 |
| 25 | 18.0000 | 18.1500 | 18.3000 |
| 26 | 18.3750 | 18.5000 | 18.5625 |
| 27 | 19.1000 | 19.2000 | 19.3000 |
| 28 | 19.5000 | 19.5000 | 19.7000 |
| 29 | 20.1500 | 20.2000 | 20.3500 |
| 30 | 20.5000 | 20.5000 | 20.5750 |
| 31 | 21.0000 | 21.2500 | 21.3750 |
| 32 | 21.5750 | 21.6500 | 21.8000 |
| 33 | 22.1000 | 22.1000 | 22.3000 |
| 34 | 22.6000 | 22.6500 | 22.8000 |
| 35 | 23.0000 | 23.1000 | 23.2500 |
| 36 | 23.2500 | 23.4500 | 23.7250 |
| 37 | 23.7500 | 23.8000 | 23.8900 |

**Table 3.** Anterior-posterior diameter percentiles

| Gestational<br>age(weeks) | 25 | 50 | 75 |
| --- | --- | --- | --- |
| 18 | 12.4000 | 12.4500 | 12.6500 |
| 19 | 12.6450 | 12.7250 | 12.8400 |
| 20 | 13.3500 | 13.4000 | 13.5500 |
| 21 | 13.7375 | 13.8000 | 13.8625 |
| 22 | 14.2325 | 14.2550 | 14.3750 |
| 23 | 14.6000 | 14.7000 | 14.7500 |
| 24 | 15.2000 | 15.2500 | 15.4000 |
| 25 | 15.6875 | 15.7450 | 15.8000 |
| 26 | 16.2000 | 16.2750 | 16.4125 |
| 27 | 16.8000 | 16.8300 | 16.8750 |
| 28 | 17.2400 | 17.2700 | 17.3000 |
| 29 | 17.6500 | 17.7000 | 17.7550 |
| 30 | 18.4075 | 18.4450 | 18.4575 |
| 31 | 18.9050 | 18.9450 | 18.9575 |
| 32 | 19.3200 | 19.3600 | 19.3850 |
| 33 | 19.7500 | 19.8000 | 19.8850 |
| 34 | 20.3000 | 20.3400 | 20.3600 |
| 35 | 20.7500 | 20.8500 | 20.8800 |
| 36 | 21.2400 | 21.3100 | 21.3350 |
| 37 | 21.7500 | 21.8500 | 21.8700 |

**Figure 12:**
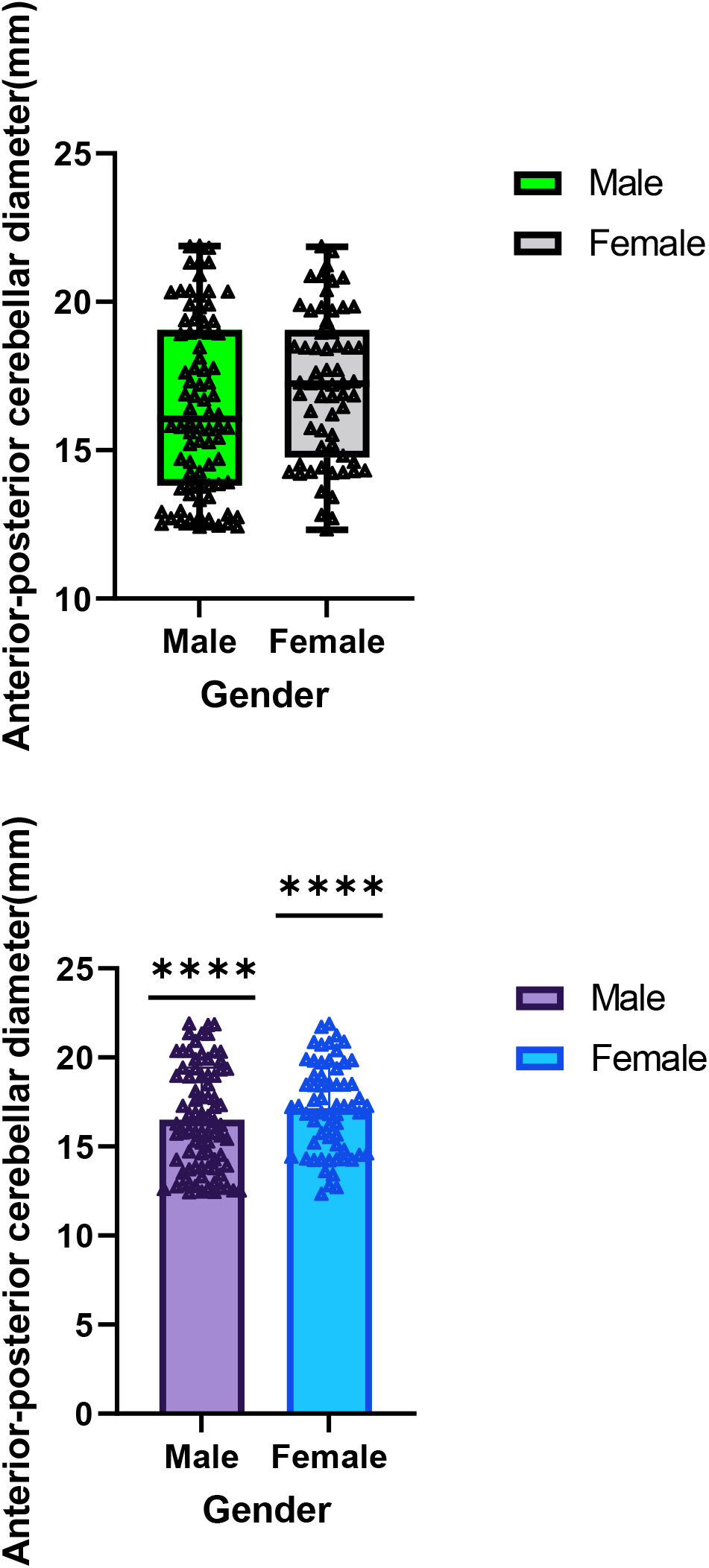
Box plot and median error pairwise comparison shows the difference in anterior-posterior between males and females

**Figure 13.**
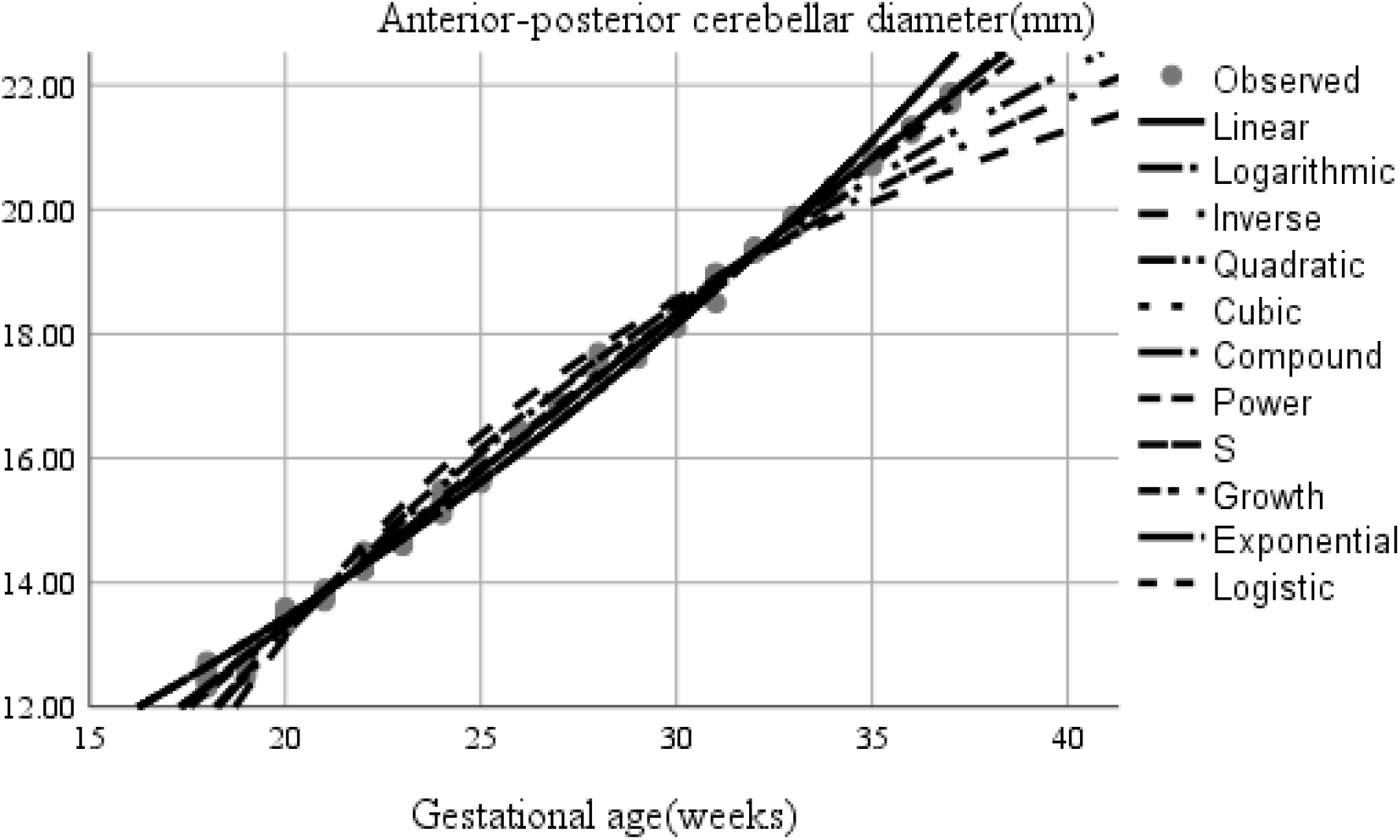
Shows several mathematical models tested, the linear being the best fit regression model

**Figure 14:**
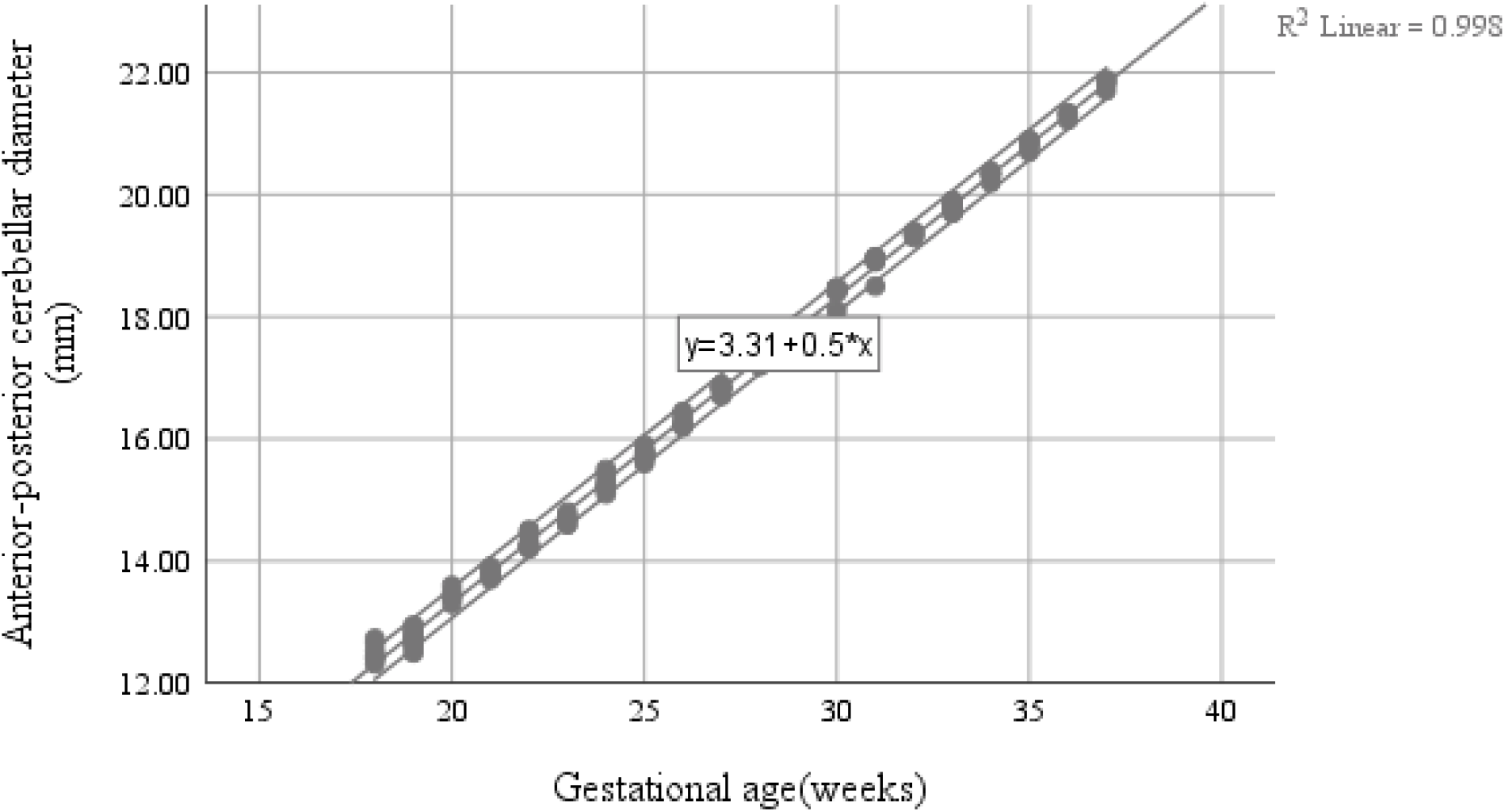
The anterior-posterior diameter and the gestational age, a linear function with 95%CI

**Figure 15.**
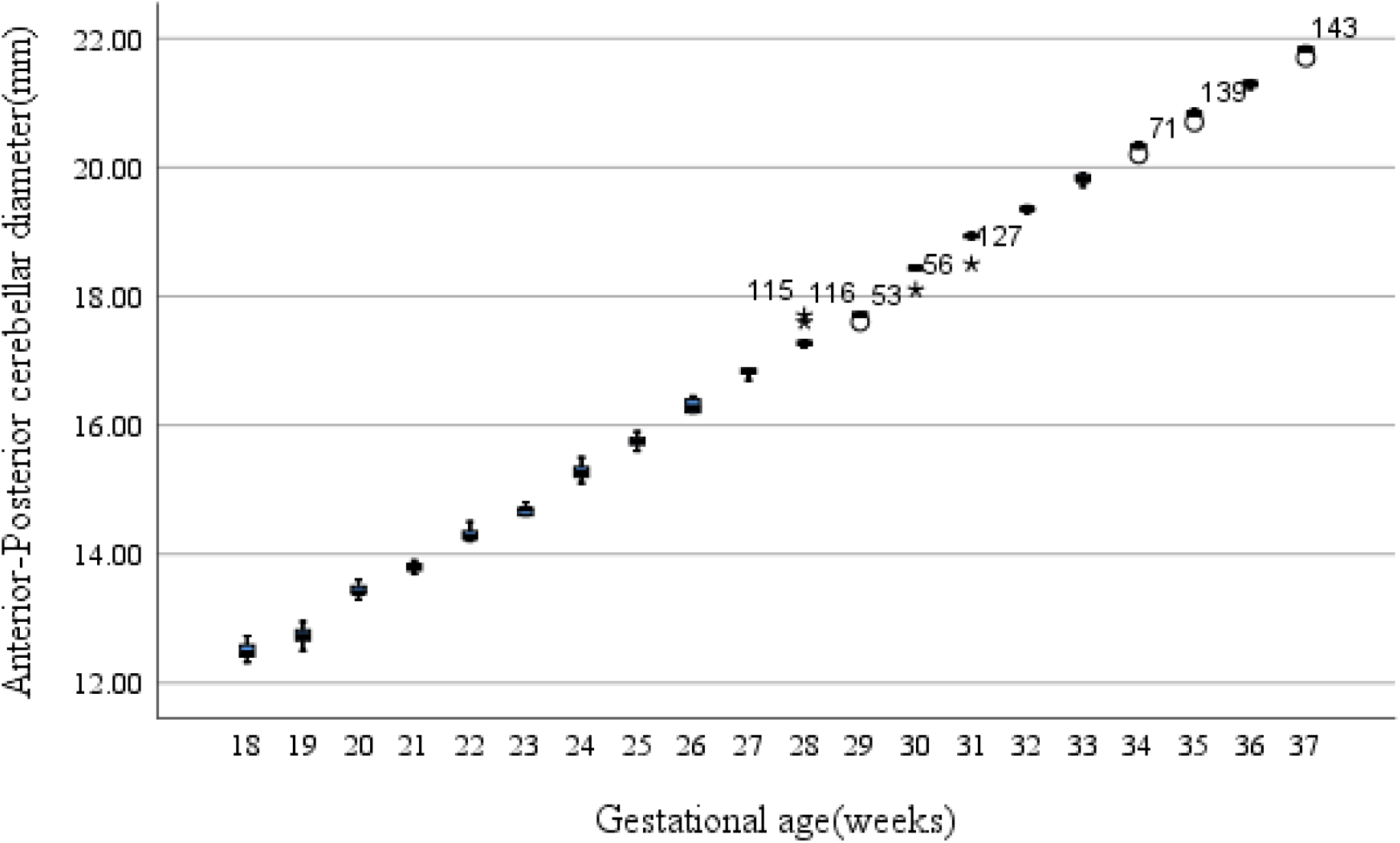
Trends of anterior-posterior cerebellar diameters and gestational age

**Figure 16.**
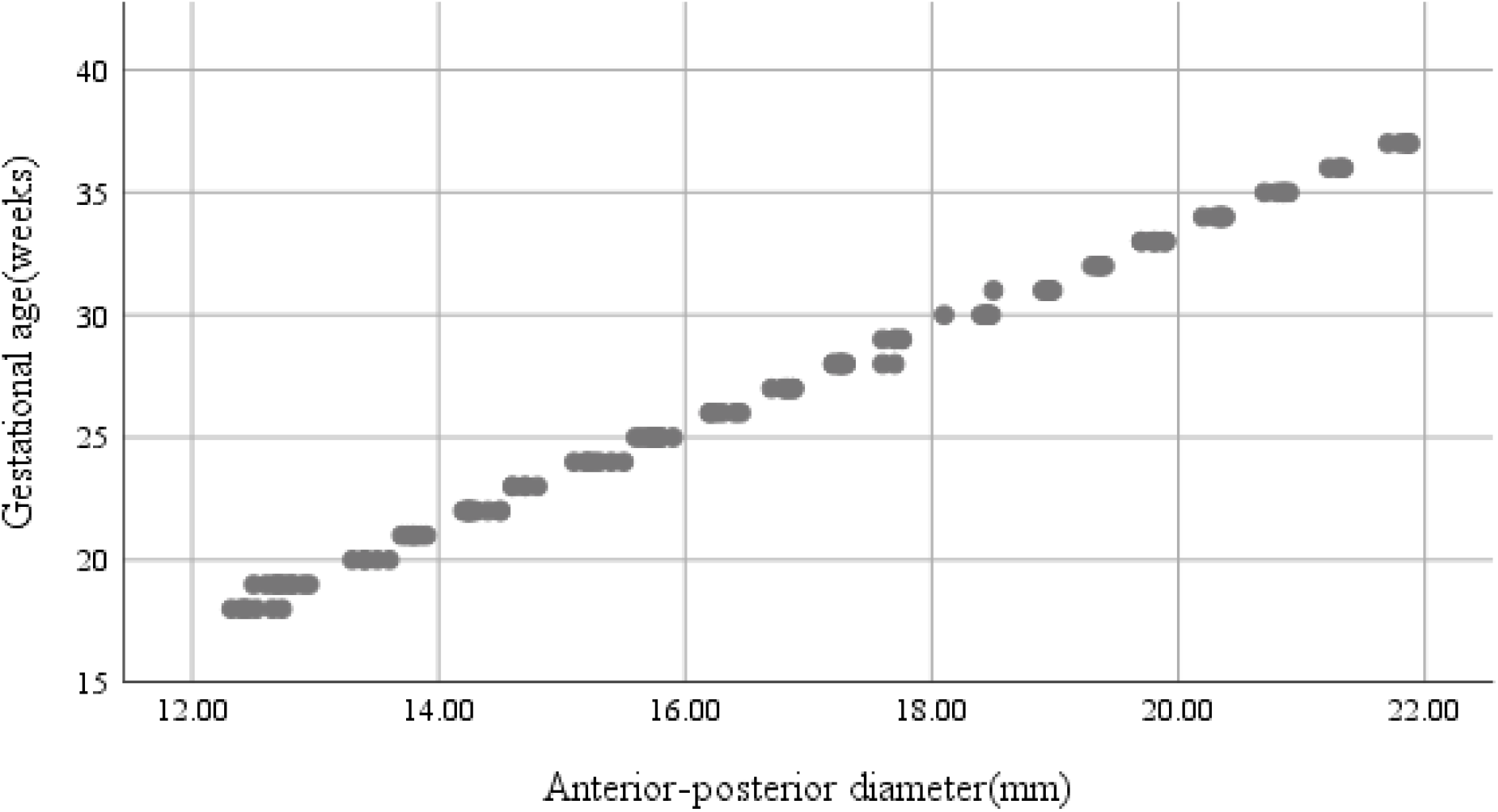
Scatter plot, anterior-posterior diameter and the gestational age

**Figure 17.**
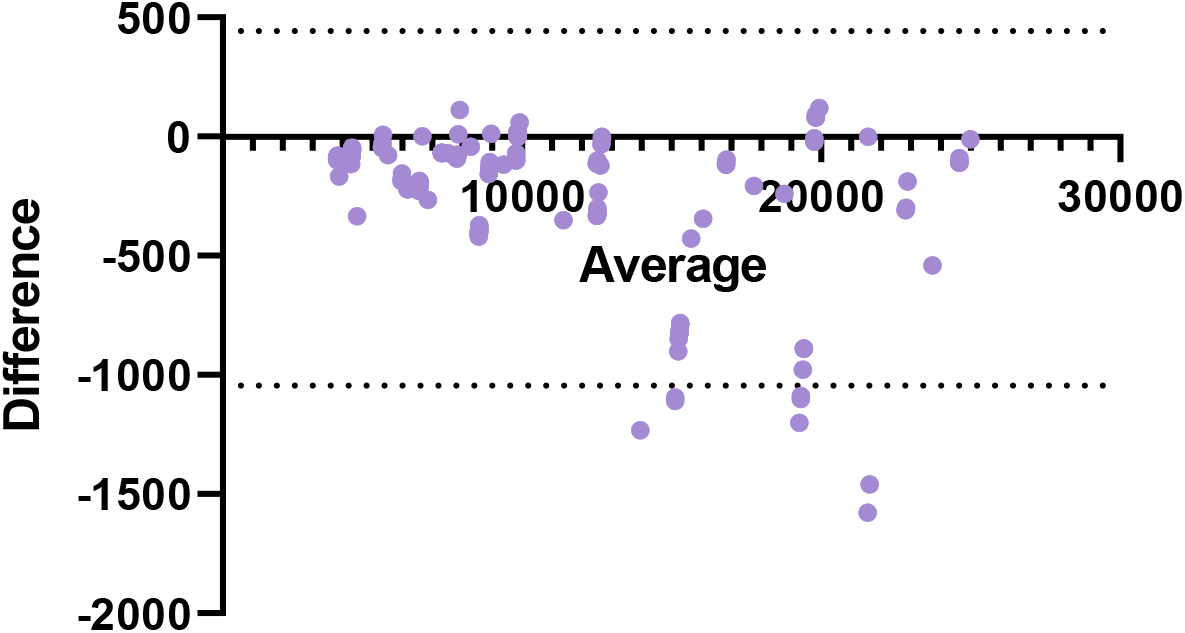
Bland-Altman plot: Difference vs. average cerebellar volume

**Figure 18.**
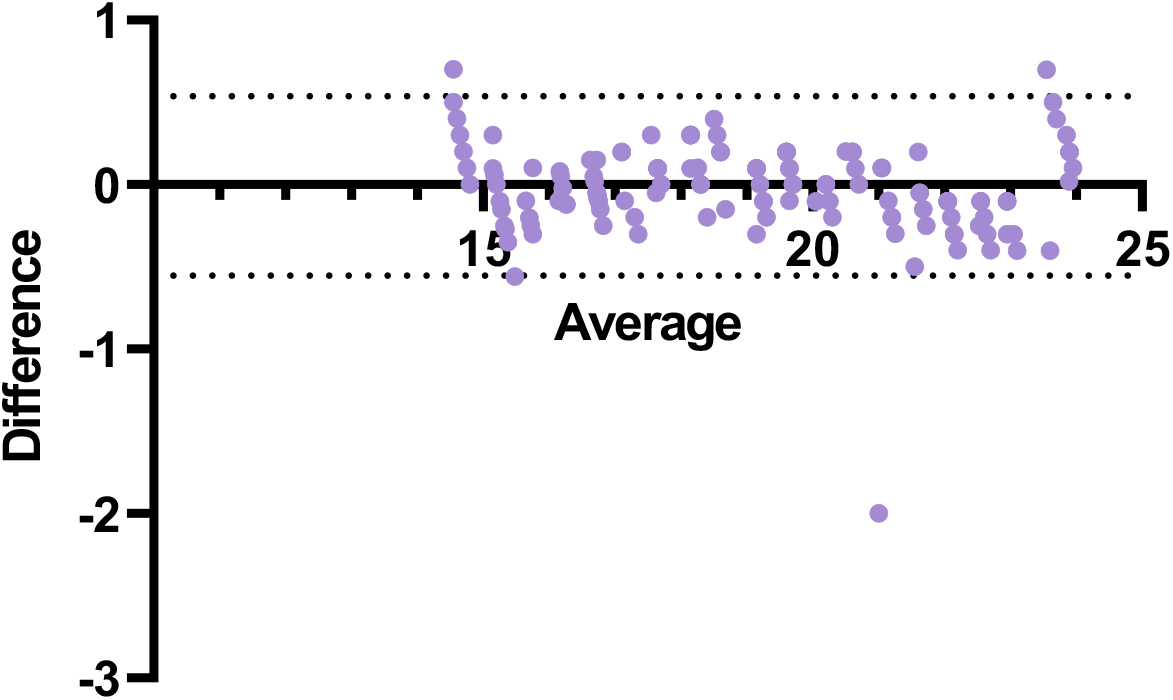
Bland-Altman plot: Difference vs. average Posterior-inferior cerebellar diameter

The median anterior- posterior diameter was 12.45 mm. The test for normality Kolmogorov-Smirnov and Shapiro-Wilk test yields the significant test of 0.22; therefore, we used Mann-Whitney test to test for sex differences and Spearman’s rho to establish the correlation between the anterior-superior diameter and the gestation age. There was significant correlation between anterior-posterior diameter and the gestational age with Spearman’s rho of (0.997, p=0.01). There was no statistically significant difference between the two groups (u=2194, p=0.16) with high mean ranks among males (78.11) as compared to females(68.26).The linear model was the best fit presenting the (y=b_o_+b_1_t) (Fig. 13 and Fig. 14) relationship between anterior-posterior diameter and the gestational age(y=3.31+0.5t) r^2^=0.998,F=70646.838. The inter-observer variability Kappa=0.6(p<0.001), 95% CI (0.306, 0.894), there was substantial agreement between the two observers in the measurements of anterior-posterior diameter. In addition, we plotted the Bland-Altman plot as shown (figure 19). The anterior- posterior diameter cerebellar diameter was increasing with the gestation age as shown in the scatter diagrams (Fig.16). The 25, 75 and 95 cerebellar volume percentiles were calculated (Fig.15) and (Table 2). The comparison between males and females are shown in box plots (Fig.12)

**Figure 19.**
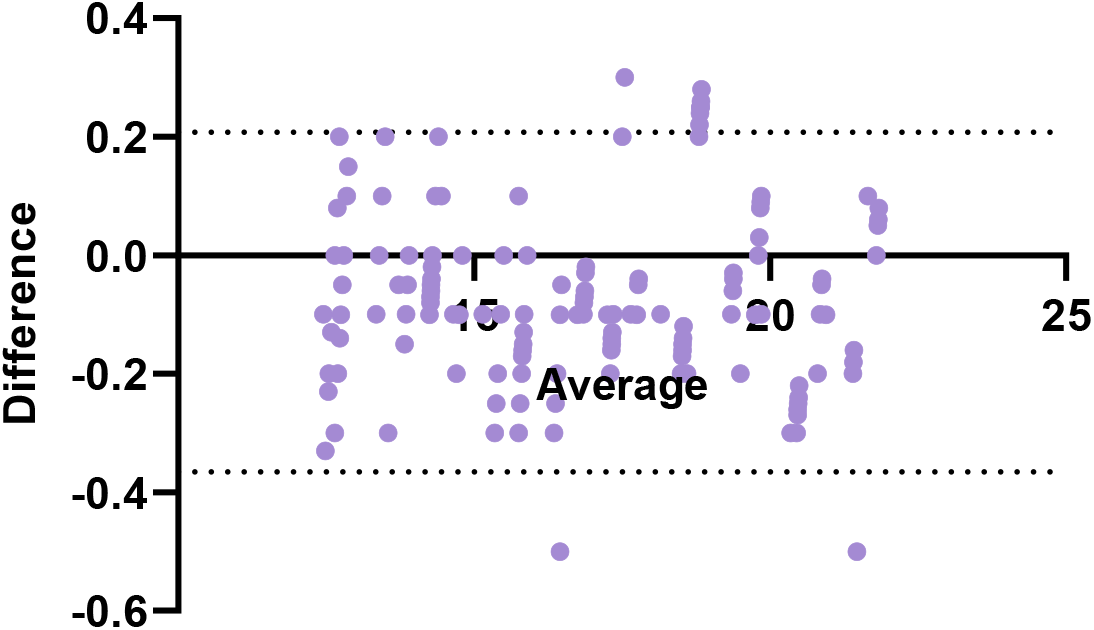
Bland-Altman plot: Difference vs. average Anterior-posterior cerebellar diameter

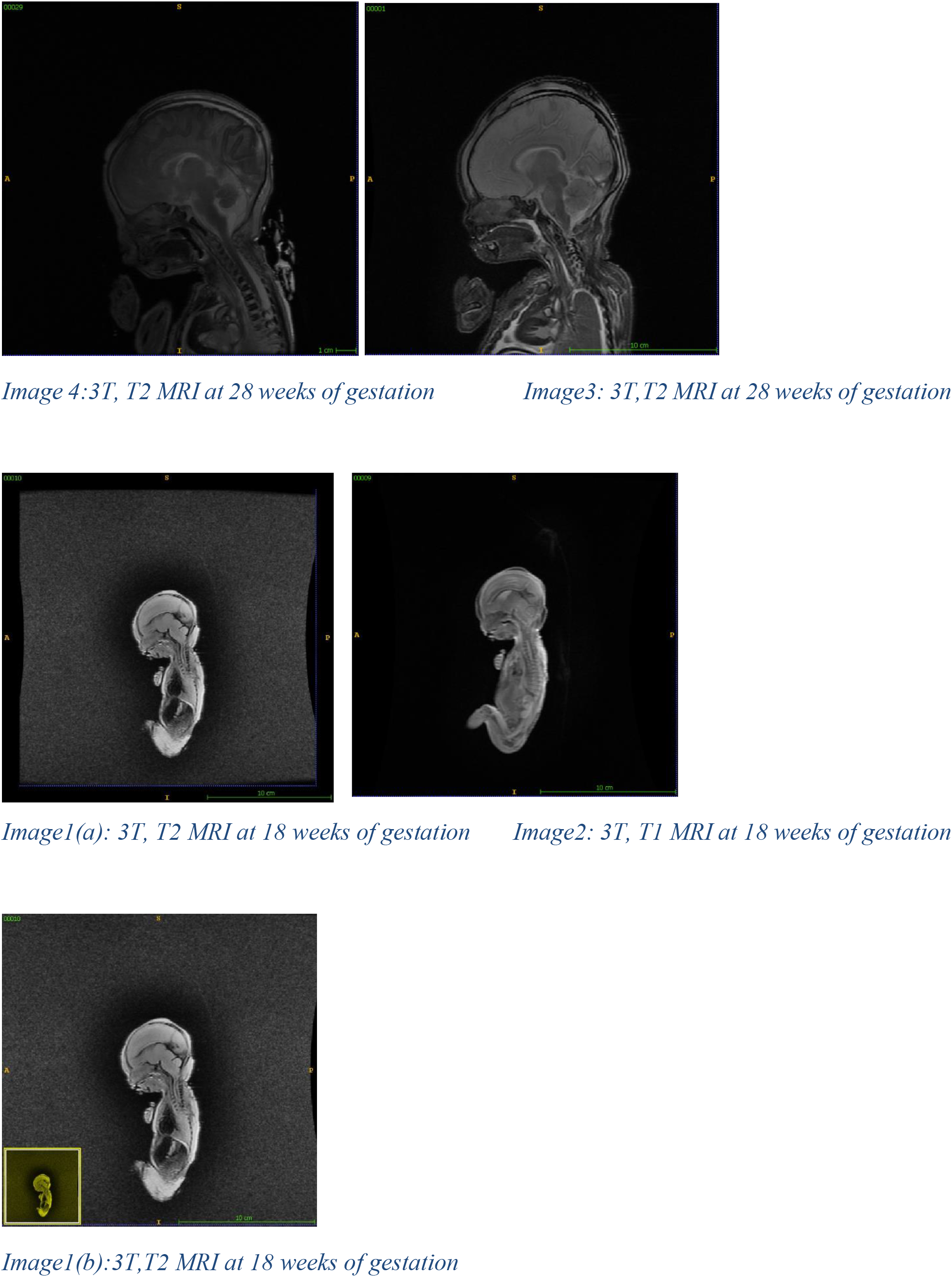
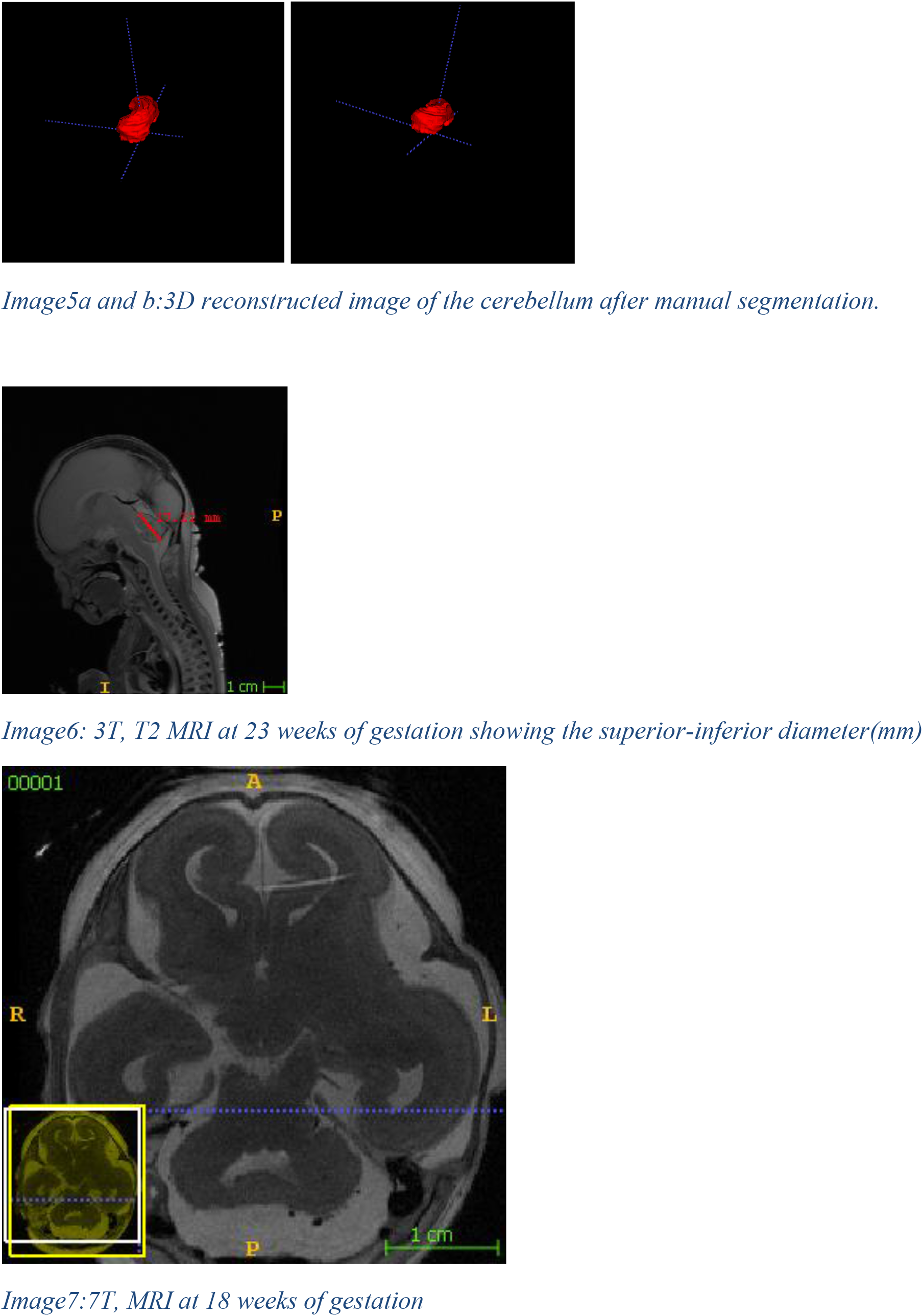

## Discussions

Magnetic resonance imaging is a valuable tool in the prenatal evaluation and can be used to supplement ultrasonography gestation imaging (Porat et al. 2002) and it is superior to the computer tomography scan because of the lack of streak artifacts and provides better contrast discrimination(Lee et al. 1984).Magnetic resonance imaging has been found to be highly accurate in the assessment of cerebellar development and the evaluation of fetal abnormalities when the ultrasonography is inconclusive(Triulzi, Parazzini, and Righini 2006) and it can also predict the neurodevelopmental outcomes (Woodward et al. 2006). The distinction between the normal and abnormal parameters of the growing cerebellum is important for establishment correct diagnosis of posterior fossa malformations(Bertucci et al. 2011).The creation of MRI template of normal cerebellar growth is very important in order to differentiate between the normal and the abnormal growth(Chong et al. 1997).The study to evaluate the normative fetal brain growth using MRI found the rate of cerebellum maturation rate was high with 4 fold increase in volume from 25-36 weeks of gestation(Clouchoux et al. 2012),this is more or less similar to our study which shows 3 fold increase in volume. In another study the cerebellar volume increased seven fold from 20 to 31 weeks of gestation with the exponential curve(R^2^=0.96) being the best fit regression equation (Scott et al. 2012), in our study, however, the increase of the cerebellar volume was the best fit by the quadratic equation or the second order polynomial (R^2^=0.994).One study used MRI to study the cerebellar volume, there was a direct correlation between the fetal cerebellar growth and the gestation age, with the second-order polynomial equation as the best fit model(Hatab, Kamourieh, and Twickler 2008),this trend is similar to our study, However dissimilar to a study among fourteen(14) human fetuses using both ultrasound and MRI, found the exponential increase in cerebellar(ellipsoid volume) during the second trimester of pregnancy(Vulturar et al. 2018). The non-linear growth curve was observed for the growing cerebellum for the fetuses aged 20-40 weeks of gestation with the differential growth between the two cerebellar hemispheres(Rutten et al. 2009).In another study using the fetal ultrasound among two hundred and thirty one(231) healthy fetuses, there was high correlation between the cerebellar volume and the gestational age, with the second degree polynomial being the best fit model(r=0.91, p<0.0001)(Chang et al. 2000a),similar to another study using 3D ultrasound which involves fifty two (52) pregnant women at gestational age of 20-32 weeks of gestation which the cerebellar volume correlated highly with the gestational age (r=0.94, p<0.001)(Araujo et al. 2007).Using 1.5T MRI among 56 fetuses of 25-41weeks of weeks of gestation, the cerebellum increase with the gestation age (r=0.51, p<0.0066(Grossman et al. 2006).In another s study to assess the fetal cerebellum, the relationship between the anterior-posterior diameter follows the first-order polynomial as the best fit predictive equation(Chang et al. 2000b) which was similar to our study. The similar study using 7T MRI scan among forty (40) Chinese fetuses, they found the correlation between the cerebellar volume the gestational age follows the second order polynomial curve in the second trimester of pregnancy (14-22) weeks of gestation(Liu et al. 2011), and another study in the same Chinese fetuses with the sample size of thirty five(35) fetuses from 15-22 weeks of gestation which shows the cerebellar volume increases by following the second- order polynomial(Xu et al. 2020),in this study, however, we used the large sample size 144 fetuses and we included the fetuses from second and third trimester of pregnancy, 18-37 weeks of gestation, we also included the 25^th^,50^th^,75^th^ for each gestational age, we showed the sex differences in cerebellar volume, superior-inferior and anterior-posterior diameters.

## Conclusion

The normative assessment of the growing cerebellum using magnetic resonance imaging is crucial for quantifying the cerebellar volume, anterior-posterior and the superior-inferior diameters. The cerebellar volume follows the second-degree polynomial while the anterior-posterior and the superior-inferior diameters follow the linear regression models. Knowing how these parameters change with the gestational age is important for accurate evaluation of the normal growing fetal cerebellum and subsequently detection of the cerebellar malformations.

## Declarations

### Ethics approval and consent to participate

Ethical clearance was provided by the University of Shandong ethical clearance committee.

### Consent for Publication

The Consent to publish this manuscript was provided by the department of Anatomy School of Basic Sciences Shandong University.

## Availability of data and material

The fetal specimens used were obtained from Qilu hospital and all the references in this manuscript were cited from PubMed and Google scholar. We used Endnote software version 8 for data management and referencing_o_

## Author’s contributions

All authors did participate in the whole process of writing this study. Emmanuel Suluba (ES), concept, drafting the manuscript, manuscript writing and reviewing the literature_o_ Liu Shuwei (LS), supervision provide guidelines, drafting, manuscript corrections and supervision_o_

## Acknowledgements

We thank the University of Shandong for providing facilities required for effective writing and submitting the manuscript.

## References

Abel, F., and M. Z. Tahir. 2019. “Role of sleep study in children with Chiari malformation and sleep disordered breathing.” Childs Nerv Syst 35 (10):1763–1768. doi: 10.1007/s00381-019-04302-0.

Anderson, P. J., J. L. Cheong, and D. K. Thompson. 2015. “The predictive validity of neonatal MRI for neurodevelopmental outcome in very preterm children.” Semin Perinatol 39 (2):147–58. doi: 10.1053/j.semperi.2015.01.008.

Araujo, Edward, Claudio Rodrigues Pires, Luciano Marcondes Machado Nardozza, Hélio Antonio Guimarães Filho, and Antonio Fernandes Moron. 2007. “Correlation of the fetal cerebellar volume with other fetal growth indices by three-dimensional ultrasound.” The Journal of Maternal-Fetal & Neonatal Medicine 20 (8):581–587. doi: 10.1080/14767050701482928.

Barkovich, A. J. 2006. “MR imaging of the neonatal brain.” Neuroimaging Clin N Am 16 (1):117-35, viii-ix. doi: 10.1016/j.nic.2005.10.003.

Basson, M. A., and R. J. Wingate. 2013. “Congenital hypoplasia of the cerebellum: developmental causes and behavioral consequences.” Front Neuroanat 7:29. doi: 10.3389/fnana.2013.00029.

Ben-Yehudah, G., S. Guediche, and J. A. Fiez. 2007. “Cerebellar contributions to verbal working memory: beyond cognitive theory.” Cerebellum 6 (3):193–201. doi: 10.1080/14734220701286195.

Bertucci, E., L. Gindes, V. Mazza, C. Re, L. Lerner-Geva, and R. Achiron. 2011. “Vermian biometric parameters in the normal and abnormal fetal posterior fossa: three-dimensional sonographic study.” J Ultrasound Med 30 (10):1403–10. doi: 10.7863/jum.2011.30.10.1403.

Chang, Chiung-Hsin, Fong-Ming Chang, Chen-Hsiang Yu, Huei-Chen Ko, and Hsi-Yao Chen. 2000a. “Assessment of fetal cerebellar volume using three-dimensional ultrasound.” Ultrasound in Medicine & Biology 26 (6):981–988. doi: https://doi.org/10.1016/S0301-5629(00)00225-8.

Chang, Chiung-Hsin, Fong-Ming Chang, Chen-Hsiang Yu, Huei-Chen Ko, and Hsi-Yao Chen. 2000b. “Three-dimensional ultrasound in the assessment of fetal cerebellar transverse and antero-posterior diameters.” Ultrasound in Medicine & Biology 26 (2):175–182. doi: https://doi.org/10.1016/S0301-5629(99)00123-4.

Chong, B. W., C. J. Babcook, D. Pang, and W. G. Ellis. 1997. “A magnetic resonance template for normal cerebellar development in the human fetus.” Neurosurgery 41 (4):924–8; discussion 928-9. doi: 10.1097/00006123-199710000-00029.

Clouchoux, Cedric, Nicolas Guizard, Alan Charles Evans, Adre Jacques du Plessis, and Catherine Limperopoulos. 2012. “Normative fetal brain growth by quantitative in vivo magnetic resonance imaging.” American Journal of Obstetrics and Gynecology 206 (2):173.e1–173.e8. doi: https://doi.org/10.1016/j.ajog.2011.10.002.

Gandolfi Colleoni, G., E. Contro, A. Carletti, T. Ghi, G. Campobasso, G. Rembouskos, G. Volpe, G. Pilu, and P. Volpe. 2012. “Prenatal diagnosis and outcome of fetal posterior fossa fluid collections.” Ultrasound Obstet Gynecol 39 (6):625–31. doi: 10.1002/uog.11071.

Grossman, Rachel, Chen Hoffman, Yael Mardor, and Anat Biegon. 2006. “Quantitative MRI measurements of human fetal brain development in utero.” NeuroImage 33 (2):463–470. doi: https://doi.org/10.1016/j.neuroimage.2006.07.005.

Haratz, K. K., S. L. Shulevitz, Z. Leibovitz, D. Lev, J. Shalev, M. Tomarkin, G. Malinger, T. Lerman-Sagie, and L. Gindes. 2019. “Fourth ventricle index: sonographic marker for severe fetal vermian dysgenesis/agenesis.” Ultrasound Obstet Gynecol 53 (3):390–395. doi: 10.1002/uog.19034.

Hatab, M. R., S. W. Kamourieh, and D. M. Twickler. 2008. “MR volume of the fetal cerebellum in relation to growth.” J Magn Reson Imaging 27 (4):840–5. doi: 10.1002/jmri.21290.

Henrichs, J., V. Verfaille, P. Jellema, L. Viester, E. Pajkrt, J. Wilschut, H. E. van der Horst, A. Franx, and A. de Jonge. 2019. “Effectiveness of routine third trimester ultrasonography to reduce adverse perinatal outcomes in low risk pregnancy (the IRIS study): nationwide, pragmatic, multicentre, stepped wedge cluster randomised trial.” Bmj 367:l5517. doi: 10.1136/bmj.l5517.

Howley, M. M., K. M. Keppler-Noreuil, C. M. Cunniff, and M. L. Browne. 2018. “Descriptive epidemiology of cerebellar hypoplasia in the National Birth Defects Prevention Study.” Birth Defects Res 110 (19):1419–1432. doi: 10.1002/bdr2.1388.

Hsu, Louis M., and Ronald Field. 2003. “Interrater Agreement Measures: Comments on Kappan, Cohen’s Kappa, Scott’s π, and Aickin’s α.” Understanding Statistics 2 (3):205–219. doi: 10.1207/S15328031US0203_03.

Imamoglu, E. Y., T. Gursoy, F. Ovali, M. Hayran, and G. Karatekin. 2013. “Nomograms of cerebellar vermis height and transverse cerebellar diameter in appropriate-for-gestational-age neonates.” Early Hum Dev 89 (12):919–23. doi: 10.1016/j.earlhumdev.2013.10.001.

Koziol, L. F., D. Budding, N. Andreasen, S. D’Arrigo, S. Bulgheroni, H. Imamizu, M. Ito, M. Manto, C. Marvel, Parker, G. Pezzulo, N. Ramnani, D. Riva, J. Schmahmann, L. Vandervert, and T. Yamazaki. 2014. “Consensus paper: the cerebellum’s role in movement and cognition.” Cerebellum 13 (1):151–77. doi: 10.1007/s12311-013-0511-x.

Lee, B. C., J. B. Kneeland, M. D. Deck, and P. T. Cahill. 1984. “Posterior fossa lesions: magnetic resonance imaging.” Radiology 153 (1):137–43. doi: 10.1148/radiology.153.1.6473775.

Liu, F., Z. Zhang, X. Lin, G. Teng, H. Meng, T. Yu, F. Fang, F. Zang, Z. Li, and S. Liu. 2011. “Development of the human fetal cerebellum in the second trimester: a post mortem magnetic resonance imaging evaluation.” J Anat 219 (5):582–8. doi: 10.1111/j.1469-7580.2011.01418.x.

Martinez, S., A. Andreu, N. Mecklenburg, and D. Echevarria. 2013. “Cellular and molecular basis of cerebellar development.” Front Neuroanat 7:18. doi: 10.3389/fnana.2013.00018.

Millen, K. J., and J. G. Gleeson. 2008. “Cerebellar development and disease.” Curr Opin Neurobiol 18 (1):12–9. doi: 10.1016/j.conb.2008.05.010.

Patek, K. J., B. M. Kline-Fath, R. J. Hopkin, V. V. Pilipenko, T. M. Crombleholme, and C. G. Spaeth. 2012. “Posterior fossa anomalies diagnosed with fetal MRI: associated anomalies and neurodevelopmental outcomes.” Prenat Diagn 32 (1):75–82. doi: 10.1002/pd.2911.

Pinchefsky, E. F., A. Accogli, M. I. Shevell, C. Saint-Martin, and M. Srour. 2019. “Developmental outcomes in children with congenital cerebellar malformations.” Dev Med Child Neurol 61 (3):350–358. doi: 10.1111/dmcn.14059.

Porat, S., R. Agid, U. Elchalal, Y. Ezra, J. M. Gomori, and M. Nadjari. 2002. “[Magnetic resonance imaging as a prenatal diagnostic tool supplementary to ultrasound in diagnosing fetal and gestational abnormalities].” Harefuah 141 (4):329–34, 412, 411.

Pugash, D., P. C. Brugger, D. Bettelheim, and D. Prayer. 2008. “Prenatal ultrasound and fetal MRI: the comparative value of each modality in prenatal diagnosis.” Eur J Radiol 68 (2):214–26. doi: 10.1016/j.ejrad.2008.06.031.

Quarello, E., M. Molho, C. Garel, A. Couture, M. P. Legac, M. L. Moutard, J. P. Bault, C. Fallet-Bianco, and Guibaud. 2014. “Prenatal abnormal features of the fourth ventricle in Joubert syndrome and related disorders.” Ultrasound Obstet Gynecol 43 (2):227–32. doi: 10.1002/uog.12567.

Rutten, M. J., L. R. Pistorius, E. J. Mulder, P. Stoutenbeek, L. S. de Vries, and G. H. Visser. 2009. “Fetal cerebellar volume and symmetry on 3-d ultrasound: volume measurement with multiplanar and vocal techniques.” Ultrasound Med Biol 35 (8):1284–9. doi: 10.1016/j.ultrasmedbio.2009.03.016.

Santoro, M., A. Coi, I. Barisic, E. Garne, M. C. Addor, J. E. H. Bergman, F. Bianchi, L. Boban, P. Braz, C. Cavero-Carbonell, M. Gatt, M. Haeusler, A. Kinsner-Ovaskainen, K. Klungsoyr, J. J. Kurinczuk, N. Lelong, K. Luyt, A. Materna-Kiryluk, O. Mokoroa, C. Mullaney, V. Nelen, A. J. Neville, M. T. O’Mahony, I. Perthus, H. Randrianaivo, J. Rankin, A. Rissmann, F. Rouget, B. Schaub, D. Tucker, D. Wellesley, L. Yevtushok, and A. Pierini. 2019. “Epidemiology of Dandy-Walker Malformation in Europe: A EUROCAT Population-Based Registry Study.” Neuroepidemiology 53 (3-4):169–179. doi: 10.1159/000501238.

Scott, J. A., K. S. Hamzelou, V. Rajagopalan, P. A. Habas, K. Kim, A. J. Barkovich, O. A. Glenn, and C. Studholme. 2012. “3D morphometric analysis of human fetal cerebellar development.” Cerebellum 11 (3):761–70. doi: 10.1007/s12311-011-0338-2.

Ten Donkelaar, H. J., and M. Lammens. 2009. “Development of the human cerebellum and its disorders.” Clin Perinatol 36 (3):513–30. doi: 10.1016/j.clp.2009.06.001.

ten Donkelaar, H. J., M. Lammens, P. Wesseling, H. O. Thijssen, and W. O. Renier. 2003. “Development and developmental disorders of the human cerebellum.” J Neurol 250 (9):1025–36. doi: 10.1007/s00415-003-0199-9.

Tonni, G., G. Grisolia, and W. Sepulveda. 2014. “Second trimester fetal neurosonography: reconstructing cerebral midline anatomy and anomalies using a novel three-dimensional ultrasound technique.” Prenat Diagn 34 (1):75–83. doi: 10.1002/pd.4258.

Triulzi, F., C. Parazzini, and A. Righini. 2006. “Magnetic resonance imaging of fetal cerebellar development.” Cerebellum 5 (3):199–205. doi: 10.1080/14734220600589210.

Vulturar, D., A. Fărcăşanu, F. Turcu, D. Boitor, and C. Crivii. 2018. “The volume of the cerebellum in the second semester of gestation.” Clujul Med 91 (2):176–180. doi: 10.15386/cjmed-922.

Warrens, Matthijs J. 2014. “New Interpretations of Cohen’s Kappa.” Journal of Mathematics 2014:203907. doi: 10.1155/2014/203907.

Whitworth, M., L. Bricker, and C. Mullan. 2015. “Ultrasound for fetal assessment in early pregnancy.” Cochrane Database Syst Rev (7):Cd007058. doi: 10.1002/14651858.CD007058.pub3.

Woodward, L. J., P. J. Anderson, N. C. Austin, K. Howard, and T. E. Inder. 2006. “Neonatal MRI to predict neurodevelopmental outcomes in preterm infants.” N Engl J Med 355 (7):685–94. doi: 10.1056/NEJMoa053792.

Wuest, A., D. Surbek, R. Wiest, C. Weisstanner, H. Bonel, M. Steinlin, L. Raio, and B. Tutschek. 2017. “Enlarged posterior fossa on prenatal imaging: differential diagnosis, associated anomalies and postnatal outcome.” Acta Obstet Gynecol Scand 96 (7):837–843. doi: 10.1111/aogs.13131.

Xu, Feifei, Xinting Ge, Yonggang Shi, Zhonghe Zhang, Yuchun Tang, Xiangtao Lin, Gaojun Teng, Fengchao Zang, Nuonan Gao, Haihong Liu, Arthur W. Toga, and Shuwei Liu. 2020. “Morphometric development of the human fetal cerebellum during the early second trimester.” NeuroImage 207:116372. doi: https://doi.org/10.1016/j.neuroimage.2019.116372.

Yang, F., T. Z. Yang, H. Luo, C. X. Li, J. Yao, X. Y. Yan, and B. Song. 2012. “[Comparative study of ultrasonography and magnetic resonance imaging in midline structures of fetal brain].” Sichuan Da Xue Xue Bao Yi Xue Ban 43 (5):720–4.

Yushkevich, P. A., Y. Gao, and G. Gerig. 2016. “ITK-SNAP: An interactive tool for semi-automatic segmentation of multi-modality biomedical images.” 2016 38th Annual International Conference of the IEEE Engineering in Medicine and Biology Society (EMBC), 16–20 Aug. 2016.

Yushkevich, Paul A., Artem Pashchinskiy, Ipek Oguz, Suyash Mohan, J. Eric Schmitt, Joel M. Stein, Dženan Zukić, Jared Vicory, Matthew McCormick, Natalie Yushkevich, Nadav Schwartz, Yang Gao, and Guido Gerig. 2019. “User-Guided Segmentation of Multi-modality Medical Imaging Datasets with ITK-SNAP.” Neuroinformatics 17 (1):83–102. doi: 10.1007/s12021-018-9385-x.

Zalel, Y., S. Yagel, R. Achiron, Z. Kivilevich, and L. Gindes. 2009. “Three-dimensional ultrasonography of the fetal vermis at 18 to 26 weeks’ gestation: time of appearance of the primary fissure.” J Ultrasound Med 28 (1):1–8. doi: 10.7863/jum.2009.28.1.1.

